# Nanoscale architecture and dynamics of Ca_V_1.3 channel clusters in cardiac myocytes revealed by single channel nanoscopy

**DOI:** 10.1101/2024.02.26.582084

**Authors:** Niko Schwenzer, Roman Tsukanov, Tobias Kohl, Samrat Basak, Izzatullo Sobitov, Fitzwilliam Seibertz, Rohan Kapoor, Niels Voigt, Jörg Enderlein, Stephan E. Lehnart

## Abstract

The clustering of L-type calcium channels for functional regulation of intracellular calcium signaling remains poorly understood. Here we applied super-resolution imaging to study Ca_V_1.3 channel clusters in human iPSC-derived atrial cardiomyocytes (hiPSC-aCM) to analyze subcellular localization, dimensions, architecture, and dynamics, which were largely unexplored previously. STimulated Emission Depletion (STED) imaging characterized the localization and structure of Ca_V_1.3 channel clusters in living cardiomyocytes. DNA Points Accumulation for Imaging in Nanoscale Topography (DNA-PAINT) achieved true molecular resolution, revealing an irregular channel distribution with substantial spacing. Single Particle Tracking (SPT) showed that channels co-diffuse in confined and stationary membrane nanodomains. The cytosolic C-terminal tail of Ca_V_1.3 by itself was found sufficient for cluster formation. In conclusion, our LTCC clustering studies demonstrate that Ca_V_1.3 channel clusters consist of mobile individual channels inside defined membrane nanodomains, in contrast to previous models of dense channel packing.

## Introduction

L-type calcium channels (LTCC) are essential for maintenance and regulation of heart contractility. In cardiomyocytes, LTCC opening is triggered by action potential depolarization and leads to rapid, transmembrane calcium influx and subsequent myofilament activation. As one of two cardiac LTCC isoforms, Ca_V_1.3 drives pacemaking and may, although unknown, contribute to contractility (*1*), supported by its selective expression in atrial cells and activation at more negative potentials compared to Ca_V_1.2 (*2*). LTCC form subdiffraction-sized clusters, which facilitate calcium release, cooperative gating and protein interactions (*3*). Previous studies on Ca_V_1.3 clustering in neurons showed that alternative splicing affects Ca_V_1.3 function and cluster formation possibly through C-terminal protein interactions with Calmodulin (CaM; *4*), PDZ-binding proteins (*5-7*) and Junctophilin isoforms (*8*). These findings point towards analogous modulatory mechanisms governing Ca_V_1.3 channel function in cardiomyocytes.

Recent studies revealed new regulatory mechanisms of the cardiac channel homolog Ca_V_1.2 (*9, 10*) and it has become clear that the ‘classic’ model of functional upregulation by direct channel phosphorylation is incorrect: β-adrenergic upregulation of Ca_V_1.2 currents is mediated by the small GTPase Rad, even when all potential phosphorylation sites on the α- and β-channel subunits have been removed (*11*). An alternative and possibly converging model of LTCC regulation involves the modulation of cooperativity via channel clustering (*10, 12*), but the mechanisms and molecular dynamics of channel clustering are not well understood. Earlier studies by the Santana group showed dimer-like bridging of channel C-termini mediated by Calmodulin (*4, 13*) and proposed a stochastic self-assembly model of cluster formation (*14*), however there is no direct experimental evidence for oligomerization-like cluster assemblies. Recently, increased clustering upon phosphorylation of the Cterminal Ca_V_1.2 residue S1928 by PKA in vascular cells was reported (*12*). Clustering of the Ca_V_1.3 isoform in cardiomyocytes was so far not characterized, as the channel is not expressed in ventricular cells and presents the challenge of combining adequate cell isolation, protein labeling and super-resolution microscopy.

In this study, we show that human induced pluripotent stem cell-derived atrial cardiomyocytes (hiPSC-aCM) expressing tagged Ca_V_1.3 channels present a valuable experimental approach for unraveling LTCC clustering mechanisms. We introduce a HaloTag-Ca_V_1.3 fusion protein for live-cell STimulated Emission Depletion (STED) imaging and Single Particle Tracking (SPT). Further, we use a corresponding GFP fusion protein to perform DNA Points Accumulation for Imaging in Nanoscale Topography (DNA*-*PAINT) at molecular-scale resolution (*15*), which was not previously reached in LTCC imaging studies. Combining the results of these super-resolution imaging techniques, we address the molecular spatial arrangement of clustered channels within membrane nanodomains, which was previously unclarified for LTCC in contrast to other ion channels. These findings serve as an important foundation for future studies aiming to correlate cluster structure and its modulation with functional readouts.

## Results

### Halo-tagged Ca_V_1.3 channels form clusters in the plasma membrane of atrial cardiomyocytes

Human induced pluripotent stem cell-derived atrial cardiomyocytes (hiPSC-aCM) were used as a model for the cellular physiology of atrial heart muscle (*16*). To investigate the clustering of Ca_V_1.3 calcium channels, an expression construct encoding the pore-forming subunit α_1D_ of human Ca_V_1.3 fused to an N-terminal HaloTag (Halo-Ca_V_1.3) was expressed in hiPSC-aCM using transient transfection (Fig. 1A, upper left). While the N-terminal tagging position was previously reported as functionally inert (*9, 17, 18*), we confirmed a physiological voltage-current response for our construct by patchclamp measurement (Fig. S1).

**Figure 1.**
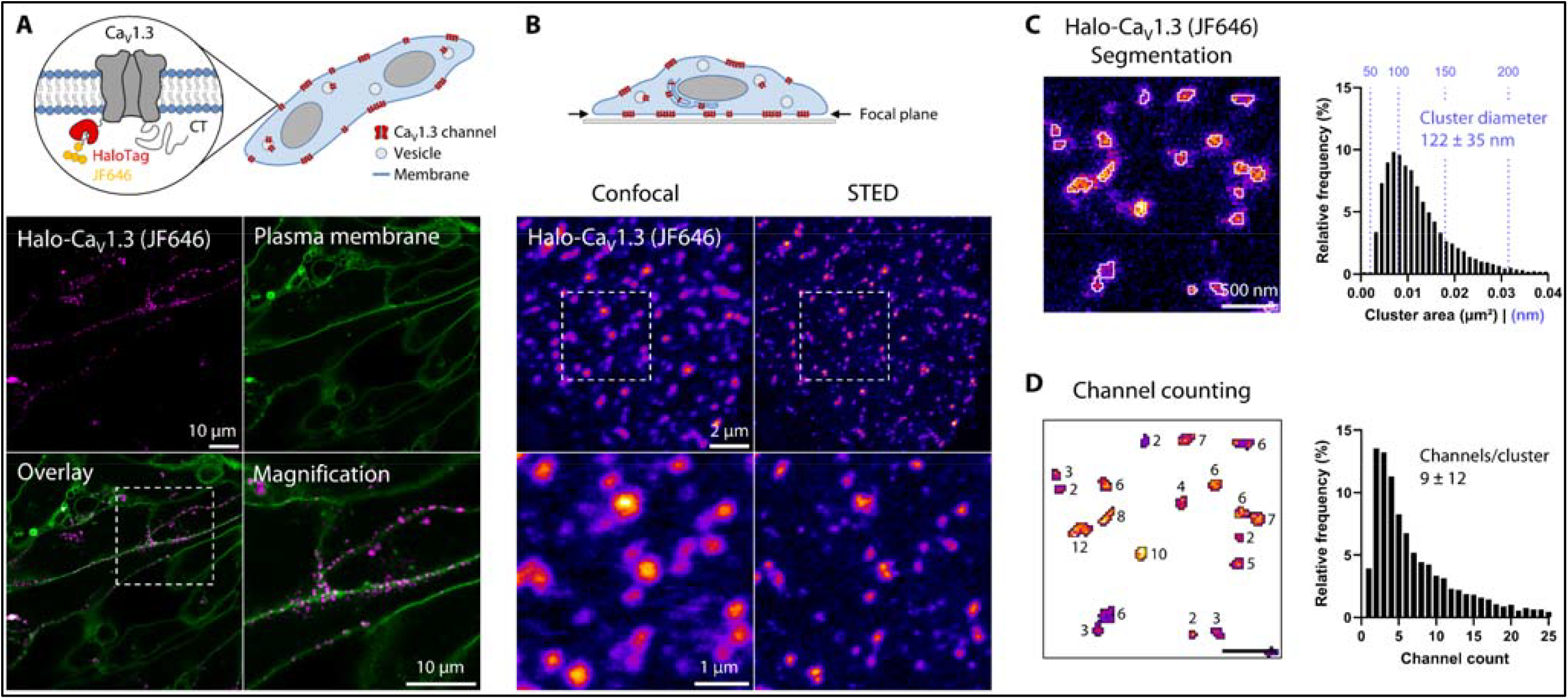
Live-cell STED imaging resolves clustering of Halo-Ca_V_1.3 at the cell surface of hiPSC-aCM. **A)** Halo-Ca_V_1.3 fusion protein was transiently expressed in hiPSC-aCM and labeled with HTL-JF646. Live-cell confocal imaging of medial cell sections revealed predominantly spot-like signal patterns of Halo-Ca_V_1.3 (magenta) distributed along the plasma membrane (green, Cholesterol-Star488), as highlighted by magnification of the indicated area. **B)** Quantitative live-cell imaging was performed centering the focal plane on the basal plasma membrane (,Fire’ LUT). STED imaging (right column) revealed sub-diffraction size and spacing of Halo-Ca_V_1.3 clusters, which could not be resolved by confocal microscopy (left). The indicated image region is magnified in the lower images, showing representative signal distributions. **C)** STED images were analyzed by automated image segmentation to detect individual signal clusters at a density of 2.0 μm^-2^ Cluster sizes averaged to areas of 0.013 ± 0.008 μm^2^, which were equivalent to diameters of 122 ± 35 nm assuming circular shape (n = 10875 clusters, N = 30 cells, 2 transfections). **D)** The signal mass of each cluster was referenced to calibration samples for molecular counting of labeled Halo-Ca_V_1.3 inside these clusters (see Fig. S2). On average clusters contained 9 ± 12 Ca_V_1.3 channels contributing to an intra-cluster channel density of 612 μm^-2^.

For live-cell imaging of hiPSC-aCM expressing Halo-Ca_V_1.3, the cells were labeled with a cell-permeable HaloTag ligand (HTL) conjugated to a fluorogenic JF646 fluorophore. Subsequent confocal imaging revealed spot-like signals (Fig. 1A, magenta) representing Ca_V_1.3 channel clusters. The signals localized predominantly to the plasma membrane, which was confirmed by co-staining with fluorescently labeled Cholesterol (green). Spot-like Ca_V_1.3 signals were only observed in transfected cells and not in apparently untransfected neighboring cells, highlighting the specificity of our labeling approach. The focal plane was then shifted to the coverslip-adherent plasma membrane of cardiomyocytes, resulting in the highest density of spot-like Halo-Ca_V_1.3 signals. To accurately resolve individual clusters below the diffraction limit, super-resolution STED imaging was applied (Fig. 1B). Compared to confocal imaging, more distinct and smaller signal shapes were detected by STED, presumably representing individual clusters. Interestingly, cluster signals often appeared in grouped arrangements, which corresponded to unresolved, single spots in the confocal image.

To quantify the abundance and size of cluster signals, image analysis was applied to a larger dataset of equivalently recorded STED images (Fig. 1C). Due to the high variability of signal spot intensities, thresholding-based methods were insufficient for segmentation and a custom approach based on robust peak finding and expansion was implemented (see Methods section). This segmentation method led to reliable cluster detection even for low-intensity or directly adjacent signals, resulting in a spatial density of 2.0 ± 0.5 (mean ± s.d.) clusters/μm^2^. Exemplary segmentation outlines are presented on raw image data in Fig. 1C. Across the dataset, these outlines encompassed cluster areas of 0.013 ± 0.008 μm^2^ corresponding to equivalent diameters of 122 ± 35 nm when assuming circular shapes. The histogram of measured areas shows a right-skewed frequency distribution, demonstrating that cluster diameters were typically around 100 nm and rarely exceeded 200 nm.

Using the same image data, molecular counting of dye molecules was applied to segmented clusters by referencing their brightness against calibration samples with defined dye numbers (Fig. S2). HaloTag labeling is well-suited for this approach, since precisely one dye molecule is covalently bound to each labeled channel. Hence the approximate number of Ca_V_1.3 channels within each cluster signal can be extracted with low statistical variance, despite not considering that a fraction of channels is unlabeled. The mean background signal surrounding each cluster signal was subtracted from the contained signal intensity. As a result, we found 9 ± 12 channels per cluster in a right-skewed distribution (median = 5, Fig. 1D). By relating the channel count of each cluster to its area, we were able to calculate intra-cluster channel densities, which amounted to 612 channels/μm^2^. Notably this result greatly differs from a theoretical limit of ∼ 10,000 channels/μm^2^ expected for oligomer-like dense channel packing, which was not nearly reached in our measurement (99^th^ percentile: 1625 channels/μm^2^).

### DNA-PAINT resolves channel arrangements and confirms loosely packed cluster structure

The molecular architecture of LTCC clusters has not been resolved thus far. Ground State Depletion (GSD) and STochastic Optical Reconstruction Microscopy (STORM) were previously used for super-resolution cluster imaging but did not reach true molecular-scale resolution. Moreover, antibody-based labeling has been a hindering factor, due to the large physical displacement (so-called linkage error) between label and epitope. We aimed to surpass these limitations by combining direct Ca_V_1.3 channel tagging with DNA-PAINT, a technique that reaches molecular-scale resolution through the use of exchangeable fluorophores – the main limiting factor in single-molecule fluorescence microscopy (*15*).

Analogous to our Halo-Ca_V_1.3 construct, we expressed GFP-Ca_V_1.3 in hiPSC-aCM and labeled for DNA-PAINT in fixed cells using the commonly used GFP nanobody (Fig. 2A; *19, 20*). Imaging was performed using a custom-built Total Internal Reflection Fluorescence (TIRF) setup with single-molecule sensitivity to image Ca_V_1.3 channels selectively in the coverslip-adherent plasma membrane. Single-molecule binding events of Atto 643 (or Atto 550)-labeled imager to its complementary docking strand were highly specific and sparsely distributed (Fig. 2B). The emitter positions were localized over time series of 30,000 to 50,000 frames to build a super-resolution image reconstruction. The resulting DNA-PAINT images showed clustered signal distributions, which were in full agreement with GFP fluorescent signals recorded at diffraction-limited resolution, confirming the specificity of DNA-PAINT binding events. A magnified image region shown in Fig. 2C demonstrates that clusters of GFP-Ca_V_1.3 were super-resolved by DNA-PAINT, leading to groups of puncta corresponding to each diffraction-limited GFP-fluorescence spot. Individual cluster magnifications (Fig. 2D) revealed a disordered arrangement of clearly separable puncta. Since puncta appearance was mostly uniform and non-overlapping, countable puncta were assumed to reflect single channel positions.

**Figure 2.**
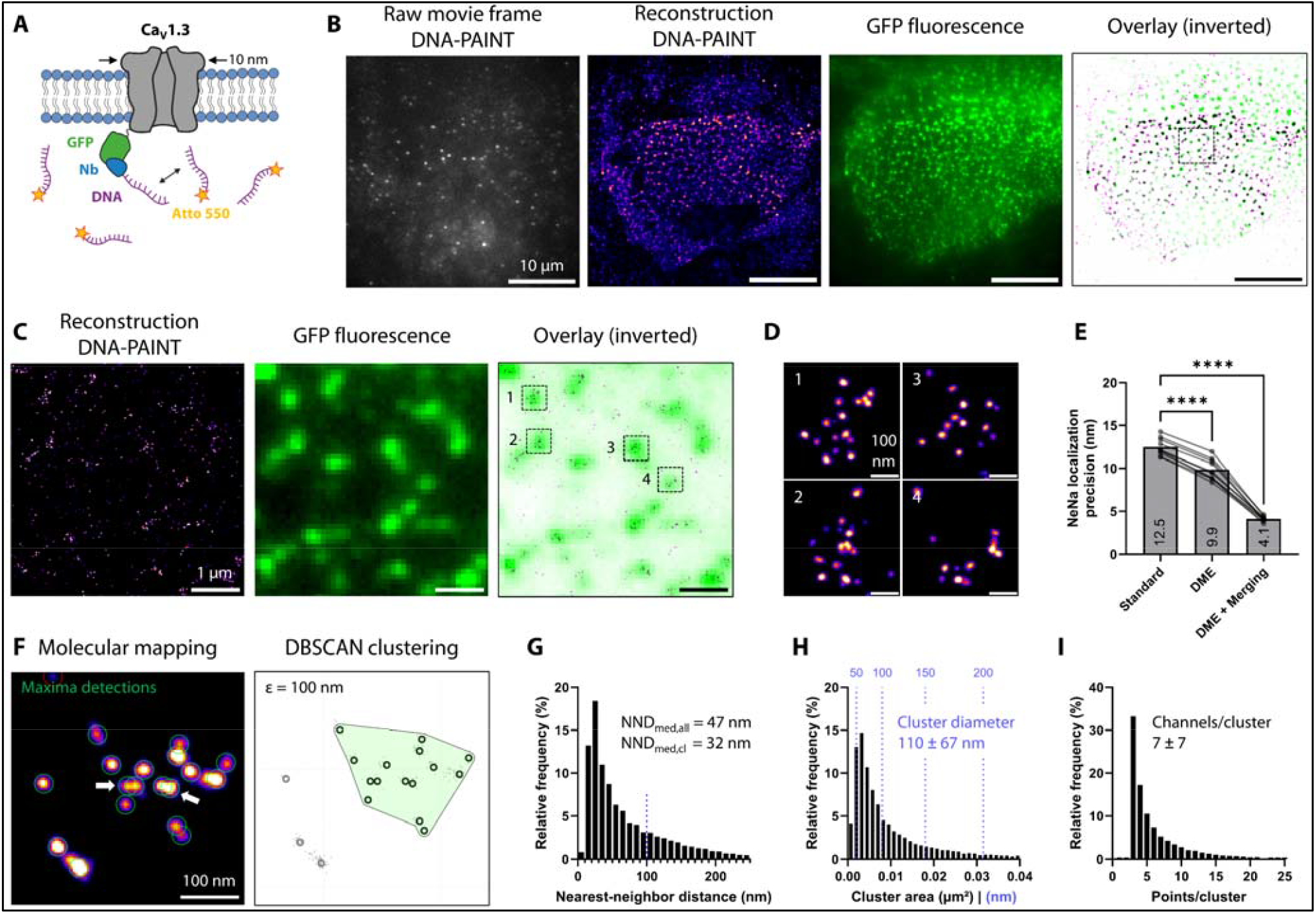
Super-resolution DNA-PAINT imaging of GFP-Ca_V_1.3 resolves channel arrangement within clusters. **A)** DNA-PAINT labeling of GFP-tagged Ca_V_1.3 expressed in hiPSC-aCM. After fixation, GFP was detected by docker-DNA-coupled anti-GFP nanobodies (NbGFP) and reversibly binding imager-DNA labeled by Atto643 or Atto550. **B)** Single raw data TIRF images of imager dye signals were used to localize DNA-PAINT binding events, and accumulated to 30,000–50,000 frames for reconstructions (,Fire’ LUT, blurred for better visibility). Simultaneous GFP fluorescence confirmed basal membrane imaging planes in GFP-Ca_V_1.3 transfected cells. **C)** Magnification of the indicated image region in (B), with clustered DNA-PAINT localization spots (left) and GFP fluorescence (center). Spatial correlation in overlay images (right) confirms specificity of anti-GFP labeling with nanobodies. **D)** Further magnifications from boxes in (C) confirm separate, countable localization spots for molecular mapping and counting of Ca_V_1.3 channel numbers. **E)** Drift correction (DME) and spot merging (binning across 4 frames) improved the NeNA-measured localization precision from 12.5 to 4.1 nm (N = 10 cells, 1 transfection). Significance was shown by repeated measures ANOVA with indicated pairwise comparisons (**** = p < 0.0001). **F)** Mapping of single channel positions was achieved by local maxima detection (green circles) from DNA-PAINT reconstructions with ≤ 5 nm localization precision (left). White arrows indicate the resolution of spots at 12 nm distance. DBSCAN clustering defined individual channel clusters, in this example 15 channels (right). DBSCAN parameters were ε = 100 nm and minPts = 3 (see Fig. S3). Molecular mapping was applied to n = 18129 clusters in N = 17 cells, 2 transfections. **G)** Nearest-neighbor distances (NND) were computed on molecular maps across all channel positions for DBSCAN-defined clusters. NND values have a plateau at 100 nm (blue dots) in line with the optimal ε parameter. **H)** Molecular maps and DBSCAN cluster outlines were used to determine cluster area and channel counts (I) distribution. Median and interquartile range (IQR) are shown in Table 1.

We benchmarked our reconstruction quality by Nearest Neighbor Analysis (NeNa; *21*) as the basis for further optimization. For initial reconstructions, a localization precision σ = 12.5 ± 1 nm was measured (Fig. 2E), which was improved to 9.8 ± 1.2 nm after applying drift and vibration correction based on a recently published algorithm (*22*). Moreover, localization merging (as described by *23*) and filtering (see Methods section) led to a drastic improvement of localization precision to 4.1 ± 0.3 nm, which was deemed sufficiently small to resolve individual Ca_V_1.3 channels with a channel diameter of 10 nm and hence similar expected minimal spacing (*24*). Along with these optimization steps, we observed a successive improvement of resolution in the reconstructions without a noticeable loss of spot detection sensitivity (Fig. S3A).

**Table 1:**
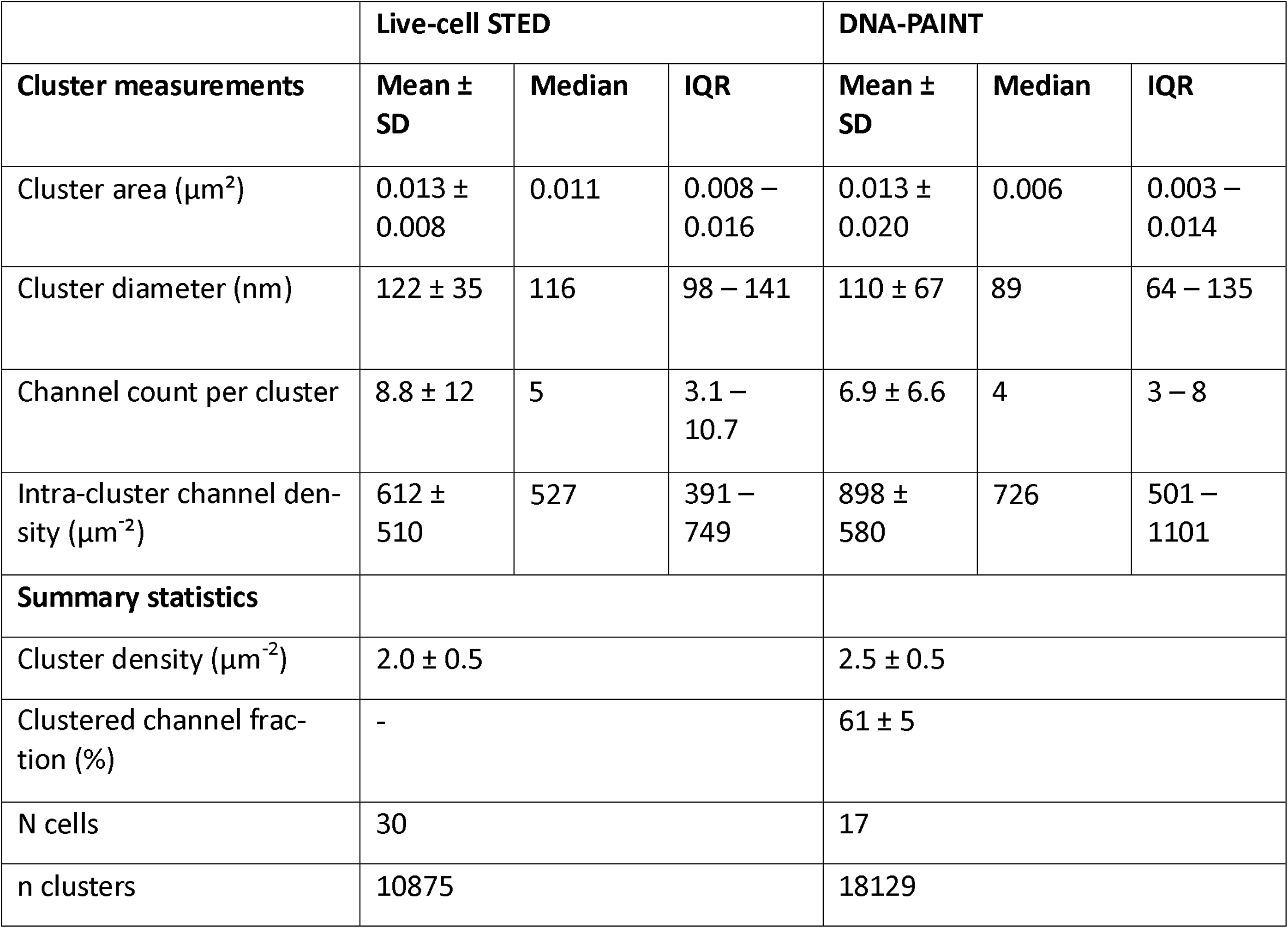
Comparison of cluster analysis results generated by STED imaging and DNA-PAINT.

To characterize clustering quantitatively, molecular mapping was performed by identification of signal maxima in DNA-PAINT reconstructions, presumably indicating single channel positions (Fig. 2F). Notably, adjacent maxima with a distance of 12 nm were reliably resolved (white arrows in Fig. 2F). Channel positions were then subjected to DBSCAN clustering (*25, 26*). The optimal parameter value ε = 100 nm was chosen and used to detect the highest number of clusters (2.5 ± 0.5 μm^-2^), whereas increased ε values led to merging of adjacent clusters and increased variance (Fig. S3B,C). We note that the detected cluster density is close to the value of 2.0 μm^-2^ obtained by STED-based cluster analysis in living cells.

Using the obtained molecular maps, we computed median nearest-neighbor distances (NND) of 47 nm and 32 nm when considering all or only clustered channels, respectively. The frequency distribution of NND for all channels (Fig. 2G) showed a local plateau at 100 nm, thus reaffirming the chosen ε value. Notably, only 19% of clustered channels were in close mutual proximity defined by NND values below 20 nm. Lastly, we quantified DBSCAN-based cluster detections, which were defined by areas of 0.013 ± 0.020 μm^2^ (Fig. 2H) containing 7 ± 7 channel spots (Fig. 2I), with both distributions showing an exponential falloff. Taken together, these results cross-validate our STED-based cluster analysis (see Table 1) and support a model of a widely spaced, disordered distribution of clustered Ca_V_1.3 channels.

### Ca_V_1.3 channels are laterally mobile despite static cluster positions

To explore the dynamics of individual Ca_V_1.3 channels within clusters, a dual labeling approach was introduced for living hiPSC-aCM expressing Halo-Ca_V_1.3: First, sparse labeling of single channels was attained by application of HTL-JF646 at minimal concentration (see Methods section). Directly after, a concurrent ensemble labeling of clusters was achieved by applying HTL-JF549 at saturating concentration. The resulting signal distributions were evaluated by live-cell, single-molecule TIRF imaging of the basal plasma membrane (Fig. 3A, left), leading to whole-cluster labeling in the JF549 channel, and alternatively well-separated single-molecule signals in the JF646 channel. For either channel, untransfected control cells showed almost no signals in comparison (Fig. 3A, center).

**Figure 3.**
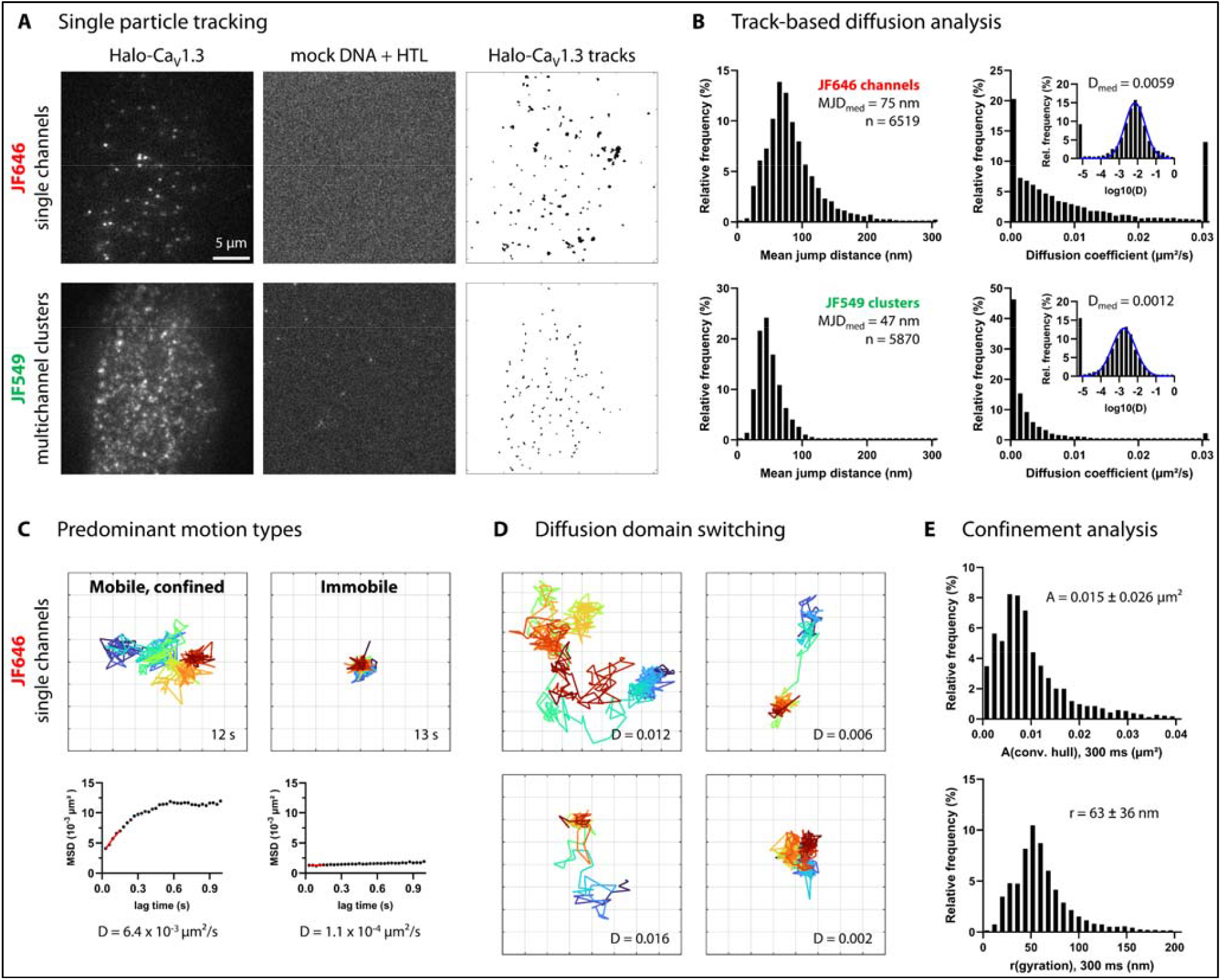
Single channel tracking quantifies the mobility and confinement of clustered Ca_V_1.3 channels. **A)** Single-molecule TIRF images of a living hiPSC-aCM expressing Halo-Ca_V_1.3 channels (first column) and a corresponding mock-transfected control cell (second column). Cells were concurrently labeled for individual channels (250 pM HTL-JF646, first row) and multichannel clusters (50 nM HTL-JF549, second row). Image series of multichannel clusters (JF549) and then single channels (HTL-JF646) were recorded consecutively. The data of both labels was independently processed by single-particle-tracking (SPT). The resulting tracks are shown as an overlay for the exemplary cell (third column). **B)** The diffusion of individual channels and clusters is compared by histograms depicting the mean jump distance of tracks and the diffusion coefficient originating from a fit of time-dependent mean-squared-displacement (MSD) curves. Insets show the same data on a logarithmic scale with gaussian fit curves in blue and median values indicated above. D value histograms include under- and overflow bins for values outside of the axis range. The dataset includes n = 6519 single channel tracks and n = 5873 cluster tracks in N = 15 cells, one transfection. **C)** Exemplary tracks of JF646-labeled single channels with duration > 10 s demonstrate the two predominant motion types: Mobile, confined (left) and immobile (right). The corresponding MSD curves are shown on the right side, with a red line indicating the linear fit used to retrieve the diffusion coefficient. The grid interval is 100 nm. **D)** Exemplary single-channel tracks show occasional domain and motion type switching, characterized by intermittently high mobility traversal between multiple domains of lower mobility and high confinement. **E)** Confinement analysis was performed on tracks containing at least 10 localizations. Local confinement was characterized as the convex hull area surrounding each complete track, and the radius of gyration. Both metrics were computed over a 300 ms (10 frame) sliding time window to avoid the inclusion of multi-domain track segments.

To evaluate single channel and cluster mobilities by single particle tracking, movies of each red (JF646) and green (JF549) channel fluorescence were recorded consecutively for each cell. The resulting particle tracks for an exemplary cell are shown as a temporal overlay (Fig. 3A, right). Notably, tracks of both labeling modes were restricted to small domains, but JF646-labeled single channel tracks occupied larger areas compared to JF549-labeled cluster tracks. In the latter case, signals originated from multiple labeled channels within subdiffraction-sized domains; therefore, an averaged, central cluster position was detected and tracked. As indicated by nearly point-like track overlays, all recorded cluster positions were strongly confined or immobile.

The mobility of individual channels and clusters was then quantified by track-based diffusion analysis (Fig. 3B). For each track, we evaluated the mean jump distance (MJD) over the 30 ms frame interval and applied mean squared displacement (MSD) analysis to retrieve the diffusion coefficient. MJD of 84 ± 43 nm were measured for single channels, showing a broad distribution of values ranging from 20 to 210 nm (98% of data). In contrast, cluster positions showed MJD of only 50 ± 19 nm, which distributed rather symmetrically ranging from 10 to 100 nm (98% of data). This indicates that individual channels diffused more rapidly and showed more heterogeneous movement compared to cluster positions. Notably, mathematical modeling of immobile positions considering localization uncertainty (see Methods section) resulted in MJD of 44 ± 12 nm, which indicates by comparison that cluster positions showed little to no mobility.

Next, we examined channel and cluster diffusion by MSD analysis. The fit of individual MSD curves along the first five lag times generated short-term diffusion coefficients (D), which are a more robust measure of diffusivity that accounts for localization error (*27*). For both single channels and cluster positions, the frequency distributions of D were approximately lognormal (Fig. 3B, right). Median D values were more than threefold higher for single channels (D = 0.0059 μm^2^/s) as compared to cluster positions (D = 0.0012 μm^2^/s), confirming a significant mobility of single channels. For reference, a threshold value of 0.001 μm^2^/s (*28-30*) is commonly used to define immobile spots, which classified 46% of clusters but only 20% of single channels as immobile based on short-term diffusion. Independently, confocal time lapse data confirmed immobility of cluster positions at lower temporal but higher spatial resolution for time scales of up to 10 minutes (Fig. S4).

### Nanodomain traversal of Ca_V_1.3 channels corresponds to dynamic channel clustering

Importantly, we validated our assumption of single-molecule labeling: We almost exclusively observed single-step bleaching in intensity traces of long tracks (Fig. S5A) and found the distribution of mean track intensity to be monomodal (Fig. S5B, top-left) reflecting that multi-labeled channels were only rarely measured and thus did not interfere with the interpretation of diffusion coefficients. We also assured that both imaging modes obtained similar spot brightness and track lengths to ensure an unbiased comparison (Fig. S5B top versus bottom).

When looking into the shape of long single channel tracks (Fig. 3C), we found two predominant motion types: First, we mostly identified mobile channels, which appeared highly confined to one or multiple membrane domains and showed variable diffusion speed. Second, with lesser abundance, we identified immobile channels, showing much smaller and consistent displacements around a defined position. For the first motion type, multi-domain diffusion was observed for particularly long tracks, with a clearly higher channel mobility across inter-domain spaces (examples shown in Fig. 3D). The observed switching of motion types across consistent nanodomains seems to reflect the occasional transit of channels from one cluster to another. Notably, the high diffusivity state generally lasted less than one second before returning to the confined state for longer time periods.

To quantify the confinement of single channels in terms of domain size, we noted that power-law fitting of MSD is unsuitable for rather short track lengths and low diffusivity compared to the localization error (*31*). Instead, we determined the convex hull area and radius of gyration for each track (Fig. 3E), which are direct geometrical measurements and thus do not rely on curve fitting (*32, 33*). We limited our analysis to a time window length of 300 ms (10 frames), which ensured consistency across variable track lengths (Fig. S4B). We thereby measured convex hull areas of 0.015 ± 0.026 μm^2^ and radii of gyration of 63 ± 36 nm for single channel tracks, which is consistent with previously determined cluster dimensions obtained by STED and DNA-PAINT (Table 1). In contrast, JF549-labeled cluster positions showed vastly smaller convex hull areas of 0.004 ± 0.003 μm^2^ and radii of gyration of 39± 16 nm, which primarily reflect the localization error around immobile positions (see Methods section).

### Ca_V_1.3 clusters robustly assemble as Ca^2+^ release units with RyR2 and Junctophilin-2

Next, we examined the organization of calcium release units (CRU) in hiPSC-aCM. CRUs are specialized membrane sites characteristic for primary cardiomyocytes, where sarcoplasmic reticulum-contained Ryanodine Receptor type 2 (RyR2) and sarcolemmal LTCC are juxtaposed as functional units, which are scaffolded by Junctophilin-2 (JPH2; *34, 35, 36*) and mediate Ca^2+^-induced Ca^2+^ release from the sarcoplasmic reticulum. While CRUs in adult atrial cardiomyocytes of highly developed species are found both at the cell surface and in intracellular tubular membrane networks (*37*), immature cardiomyocytes including hiPSC-aCM cultured 2D do not feature these membrane networks and thus inherently rely on cell-surface localized domains for calcium release. This suits well to quantitative imaging of the coverslip attached membrane in a consistent, reproducible focal plane (compare Fig. 1).

Accordingly, three-channel confocal immunofluorescence of hiPSC-aCM labeled for Halo-Ca_V_1.3, JPH2 and RyR2 showed spot-like signals for each target protein at the cell surface (Fig. 4A). All three proteins colocalized to a large degree, which resulted in white spot coloring in the overlay image. Colocalization was confirmed in the basal imaging plane (Fig. 4B), which rendered the lateral distribution of each protein across the cell surface and revealed a high number of three-channel-colocalized spots. The pattern of spatial correlation was exemplified by intensity line profiles (Fig. 4A-B bottom), showing that indeed most signal peaks constituted all three proteins. To quantify the recruitment of Ca_V_1.3 to calcium release units (CRUs), three-channel image segmentation (Fig. 4C) and colocalization analysis (Fig. 4D) were applied to a dataset of similarly recorded cell-surface images: We first analyzed the fraction of colocalized Ca_V_1.3 signal and thus found that 65% of Ca_V_1.3 signal mass was RyR2-colocalized, 61% was JPH2-colocalized, and 51% was double-colocalized, indicating an efficient channel recruitment to CRUs. The results were reproduced by inversion of secondary antibodies and imaging channels for RyR2 and JPH2 in the same experiment (Fig. 4D). Lastly, we analyzed Ca_V_1.3-colocalization from the perspective of CRU composition, defining CRU signal as the product of JPH2 and RyR2 signal. Thereby we found that 86% ± 3% of CRU signal mass colocalized with Ca_V_1.3, indicating that atrial CRUs very consistently harbor Ca_V_1.3 channels.

**Figure 4.**
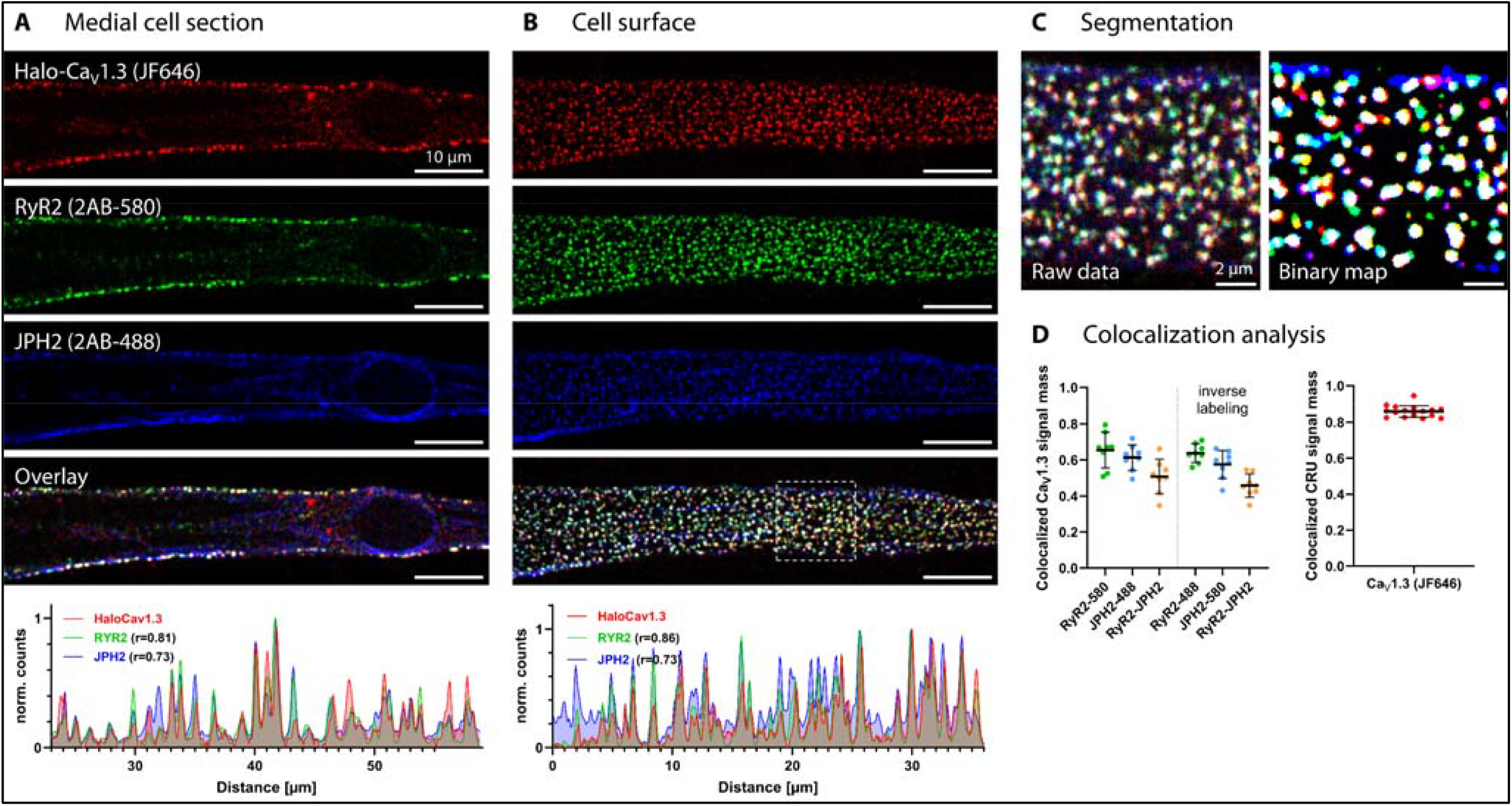
Colocalization of Ca_V_1.3 with JPH2 and RyR2 in surface-localized calcium release units. **A)** Live-cell labeling of Halo-Ca_V_1.3 expressed in hiPSC-aCM was combined with subsequent RyR2 and JPH2 immunofluorescence for confocal imaging. Medial confocal sections display Ca_V_1.3 (red), RyR2 (green) and JPH2 (blue) only in the cell periphery, where they show extensive colocalization (white coloring), representing calcium release units (CRU) localized to the plasma membrane. ‘2AB-’ indicates the secondary antibody conjugate and corresponding imaging channel (580 = Abberior Star580, 488 = Abberior StarGreen). A representative line profile across cluster signals demonstrate extensive spatial correlation between fluorescent signals corresponding to all three analyzed CRU proteins. The Pearson correlation coefficient (r) indicates one-dimensional correlation of each RyR2 and JPH2 to Halo-Ca_V_1.3 signal. **B)** Imaging of the adherent, basal plasma membrane in the same cell reveals a homogeneous 2D distribution of spot-like signals, corresponding to plasma membrane resident CRUs, as confirmed by a representative line profile. **C)** Three channel images from planar membranes (magnification from B) were segmented for signal-spots and binarized. Consequently, white signal color indicates three-channel colocalization, magenta indicates Ca_V_1.3-JPH2, yellow indicates Ca_V_1.3-RyR2 and cyan indicates RyR2-JPH2 colocalization, respectively. **D)** Colocalization analysis quantified the fraction of Ca_V_1.3 signal mass overlapping with binarized areas of either RyR2, JPH2 or both (left graph). Specific colocalization was confirmed with an inversion of fluorophores on secondary antibodies. The right graph shows the fraction of CRU signal mass (defined by the product of RyR2 and JPH2 signal) that is colocalized with Ca_V_1.3-binarized area (N = 16 cells, one transfection).

We also determined the colocalization of Halo- and GFP-tagged Ca_V_1.3 with several other subcellular compartments and proteins (Fig. S6). Live-cell co-staining with fluorescently labeled Cholesterol as a nanodomain marker showed an exclusion-like pattern rather than colocalization in the basal plasma membrane, highlighting that Ca_V_1.3 clusters are independent of cholesterol-containing lipid domains (Fig. S6A). Immunodetection of endogenous Caveolin-3 (Cav3), a marker of caveolar nanodomains, showed only mild colocalization with Ca_V_1.3 (Fig. S6B). Labeling of the endoplasmic reticulum showed no colocalization (Fig. S6C), indicating that GFP-Ca_V_1.3 was efficiently expressed and transported to the plasma membrane. No colocalization between Halo-Ca_V_1.3 and either sarcomeric alpha-actinin or Junctophilin-1 was observed (Fig. S6D,E). To test whether the association of Ca_V_1.3 and JPH2 requires other cardiac proteins, both proteins were expressed in HEK293 cells and found to strongly colocalize in the basal plasma membrane, pointing towards a tissue-independent, intrinsic association of both proteins.

### The C-terminal cytosolic tail of Ca_V_1.3 is sufficient for cluster formation

Alternative splicing of human Ca_V_1.3 primarily truncates the C-terminal protein sequence, leading to a shortening of the cytosolic C-terminal tail (CTT) from 694 to 180 amino acids in the short (42A) isoform. Since the CTT contains several important protein interaction sites (e.g. for CaM, JPH2, AKAP) and its splicing was shown to modulate channel clustering (*4, 5*), we hypothesized that the CTT may be involved in, necessary, or even sufficient to confer clustering of Ca_V_1.3 channels. We conceived an accessible experimental approach to address this by generating novel fusion proteins concatenating extracellular N-terminal HaloTag (*38*) with intracellular CTT sequence of either Ca_V_1.3 isoform. We then expressed these constructs in hiPSC-aCM and applied surface-selective labeling using HTL-Alexa488 for live-cell confocal imaging to observe the extent of cluster formation.

For both constructs and a control construct lacking CTT sequence, signals were predominantly found at the cell surface, as confirmed by co-staining of the plasma membrane (Fig. 5). Interestingly, basal plane imaging showed a segregation of signals into cluster-like shapes for both CTT constructs (Fig. 5A, magenta), but an increased abundance of clustered signals for the long versus short CTT isoform (Fig. 5B). As expected, the control construct lacking Ca_V_1.3 sequence showed a highly homogenous membrane signal (Fig. 5C), indicating that there is no intrinsic ability of the extracellular HaloTag to form clusters. The clustering of these constructs was quantified by image analysis, which confirmed a significantly higher cluster abundance for the long versus short CTT construct (mean 17.4% versus 7.9%, p < 0.0001, Fig. 5D). Notably, the surface expression level (Fig. 5E) and relative cluster brightness indicative of cluster size (Fig. 5F) were not significantly different, reflecting that CTT truncation mainly affected cluster abundance but not other parameters. The control construct of transmembrane-anchored HaloTag without CTT notably showed a much higher expression level, but a near-absence of clustering.

**Figure 5.**
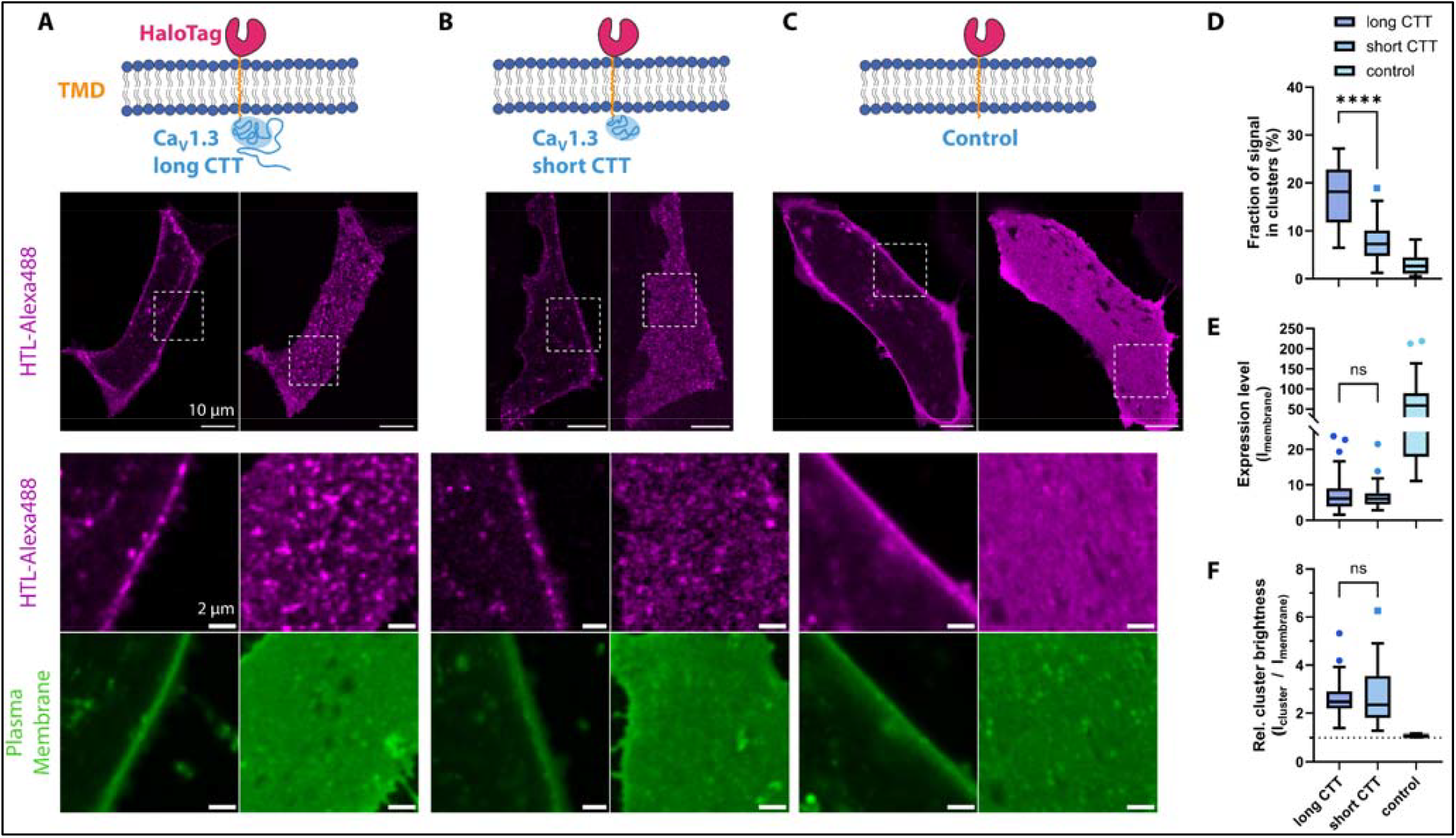
Ca_V_1.3 C-terminal construct expression in hiPSC-aCM leads to cluster formation. **A)** A fusion protein of Ca_V_1.3_42_ C-terminal tail (long CTT) attached to transmembrane-HaloTag was expressed in hiPSC-aCM using transient transfection. After live-cell labeling with cell-impermeable HTL-Alexa488, distinct spot-like signals resembling Ca_V_1.3-like clusters were revealed at the cell surface by confocal imaging in the medial (left) and basal (right) imaging plane (displayed in magenta). The localization was confirmed by co-staining with the plasma membrane marker Cellmask-DeepRed (displayed in green). **B)** Expression of the equivalent fusion protein containing the short CTT of Ca_V_1.3 splice variant 42A also lead to cluster-like signal shapes at the cell surface. However, cluster-like spots appeared less abundant compared to the long isoform shown in (A). **C)** A control construct containing only transmembrane-HaloTag showed a homogenous protein distribution in the plasma membrane without the formation of distinct, cluster-like spots. **D)** Evaluation of confocal HTL-Alexa488 signal distributions shown in (A-C) by cluster analysis. Cluster signals were detected by automatic thresholding after gaussian filtering of images and then quantified by size and brightness. Box plots show Median, IQR and Tukey-based whiskers. The fraction of clustered signal within basal plasma membranes was significantly larger for long vs short CTT (**** = p < 0.0001, Welch’s unpaired t-test, N = 28, 29, 25 cells, 2 transfections). The control condition showed an absence of significant clustering and was therefore not statistically compared. **E)** Membrane expression levels measured by the average fluorescence intensity showed no significant difference between long and short CTT (p = 0.47). **F)** Similarly, the relative signal intensity of clusters compared to the whole plasma membrane showed no significant difference between long and short CTT (p = 0.7).

Since the observed clustering of CTT constructs may depend on pre-existing endogenous Ca_V_ channel clusters in hiPSC-aCM, we similarly investigated CTT cluster formation in transfected HEK293 CT6232 cells (expressing only the accessory channel subunits β_3_ and α_2_δ_1_, but not α_1D_). In these cells lacking cardiac proteins, we again observed robust cluster formation of both CTT constructs as compared to the control construct (Fig. S7). Similar to hiPSC-aCM, a higher cluster abundance for the long versus short CTT isoform was observed. These results confirmed that Ca_V_1.3 CTT is sufficient to form Ca_V_1.3 clusters even in the absence of cardiac-specific accessory proteins and pre-existing clusters.

## Discussion

Herein nanoscale imaging of human Ca_V_1.3 channel clusters was pioneered in the hiPSC-aCM expression system with labeling strategies novel to L-type calcium channels. Consequently, we provide the first model of Ca_v_1.3 cluster assembly in cardiomyocytes, which can be readily compared to existing data in neuronal model systems. The use of hiPSC-aCM provided a physiological cellular framework for the assembly of functional calcium channels, resembling spontaneously contracting cardiomyocytes in an early developmental stage (*39, 40*). In contrast to primary cardiomyocytes, hiPSC-CM are amenable to plasmid-based gene transfection (*41*), which was harnessed in our study to transiently transfect with tagged Ca_v_1.3 variants. Channel activation was similar for wild-type and all tagged channels, while a minor change might exist for inactivation kinetics of Halo-tagged channels. Yet imaging experiments were performed at resting membrane potentials, representing channel behavior under unstimulated baseline conditions.

The atrial subtype-directed differentiation implicates an endogenous expression of LTCC in these cells (*16, 42*). Since we selectively labeled these tagged Ca_V_1.3 variants and not endogenous Ca_V_1.3 in our imaging experiments, simultaneous occurrence of both types of channels within clusters cannot be completely excluded. Offsetting this, we can assume that relative protein cell surface abundance was clearly weighted towards tagged Ca_V_1.3 due to a titration effect: in contrast to endogenous channels, tagged Ca_v_1.3 was overexpressed, while LTCC surface trafficking and residency depends on molecular assembly with β-subunits, limited by the available endogenous pool (*5, 43*). In addition, ER-resident tagged Ca_v_1.3 channels and aggregates were hardly observed in transfected cells, owing to a lower protein biosynthesis rate compared to heterologous expression systems and the sensitive unfolded protein response for ion channels in hiPSC-CM (*44*). Consequently, potential overexpression artifacts on cluster analysis are considered non-significant.

Cluster analysis was performed on the canonical, full-length human Ca_V_1.3_42_ sequence, the most abundant isoform in human cardiomyocytes (*45*). The pore-forming subunit α_1D_ was tagged at the N-terminus preserving channel voltage-gating, while the C-terminus is crucial for regulatory functions that may be perturbed by fusion-tagging. In addition, we considered that alternative tagging of the accessory β-subunit (*9, 10, 43*) was unsuitable for our study since Ca_V_β can bind other Ca^2+^ channel isoforms and performs intracellular functions (*46*). We reason that Ca_V_1.3 overexpression may have diminished the abundance of endogenous Ca_V_1.2 within the plasma membrane based on limitations of endogenous Ca_V_β subunit-dependent trafficking, therefore expecting to quantify mostly homooligomeric Halo-Ca_V_1.3 clusters, instead of endogenous Ca_V_1.2 or mixed heteromeric clusters. However, it is unclear if endogenous Ca_V_1.3 contributes to contractile activation in mature primary aCM.

We developed independent and synergistic cluster analysis workflows on hiPSC-aCM expressing tagged Ca_V_1.3 proteins. Live-cell quantitative STED and DNA-PAINT imaging determined cluster size and geometry data, while live-cell SPT produced the first mobility data on Ca_v_1.3 channels in cardiomyocytes. Both STED and DNA-PAINT detected cell surface channel clusters of on average ∼ 120 nm diameter containing 7–9 channel molecules. Notably live-cell STED excluded potential fixation artifacts previously reported (*47, 48*) and introduced channel counting based on brightness referencing (*49*). Brightness calibration assumed a similar labeling efficiency and linear signal to dye number relation for calibration beads and within cluster nanodomains in situ. The method offers a higher counting range and live-cell compatibility in contrast to widely used photobleaching analysis (*50*). Importantly, spatial fluorophore densities observed in this study vastly argue against quenching effects known for directly adjacent fluorophores (*51*).

While STED-based size metrics were limited by a spatial resolution of ∼ 70 nm, our DNA-PAINT imaging approach using GFP-targeted Ca_V_1.3 (Fig. 2) attained molecular-scale resolution down to 4 nm localization precision, which was not achieved by any previous study of LTCC clustering, e.g. compared to 16 nm in GSD imaging (*4*). We note that additional, variable displacement errors caused by the undetermined mobility of the cytosolic Ca_V_1.3 N-terminus and the physical size of the nanobody (∼4 nm) used for detection possibly reduced the effective resolution. However, these effects were deemed insignificant compared to previous approaches using indirect immunodetection (*20*). The developed procedures for DNA-PAINT and subsequent data analysis thus are sufficient to resolve adjacent channels of 10 nm diameter (*24*) and resulted in similar cluster metrics as live-cell STED imaging, confirming the validity of our approach (Table 1).

DNA-PAINT revealed a rather large, non-uniform spacing of clustered channels (median NND = 32 nm) and a low incidence of directly adjacent channels (19% of channels with NND < 20 nm). These observations argue against both oligomerization-like isotropic packing of channels and constitutive dimerization. Interestingly, we did not observe any grid-like arrangements, which are characteristic for skeletal muscle Ca_V_1.1 and RyR1 clusters (*52*). In this regard, we found that Ca_V_1.3 channel arrangement rather corresponds to the stochastic nature of cardiac RyR2 channel clusters (*53, 54*). Interestingly, a previous study in primary hippocampal neurons measured similar Ca_V_1.3 channel counts per cluster by bleach step counting and the same exponential distribution of cluster areas (*4*), although we measured approximately threefold larger cluster areas on average in cardiomyocytes. Tissue-dependent characteristics were previously not evidenced between neurons and cardiomyocytes, but rather for cochlear inner hair cells, which contain multifold larger Ca_V_1.3 clusters (*55*). Differences in cluster size could rather arise from different segmentation strategies, especially since comparison of the presented images indicates similar dimensions of DNA-PAINT and GSD-imaged clusters.

Additional data for our Ca_v_1.3 clustering model was gathered by SPT analysis of channel mobility (Fig. 3), that included both sparse single-channel labeling and ensemble labeling for tracking whole clusters. HaloTag-based covalent labeling enabled the use of bright and photostable organic fluorophores, achieving robust motion tracking by state-of-the-art algorithms (*56*). The resulting trajectories were suitable for track-based diffusion analysis and readily comparable between both imaging modes, owing to matched spot intensities and tracking durations (Fig. S5). A direct comparison revealed that individual channels exhibited much higher diffusivity than independently tracked cluster positions (median D 0.0059 versus 0.0012 μm^2^/s, respectively), thus excluding the possibility of rigidly packed cluster structures. Notably, the absence of multi-step bleaching events in our SPT data indicates that most tracks correspond to single fluorophores, and thus single channels.

Given that we identified confined mobility as the major motion type across all single channel SPT tracks, we interpreted the occupied nanodomains as being equivalent to clusters. These cluster domains were described by an average gyration radius across all single channel tracks of 64 nm, which fits to cluster diameters of around 120 nm reported by STED imaging and DNA-PAINT. Thus, our multifaceted methodological approach is devoid of major technical compromises and shows consistent independent readouts and output parameters, building up a new model of Ca_v_1.3 cluster configurations. Our SPT approach measured channel mobilities that were also in line with previously reported values of neuronal Ca_V_1.2 channels (D = 0.005 μm^2^/s; *30*). Similar to the cited study, we evidenced the traversal of channels across multiple confined domains, which implies that clusters might dynamically recruit and disband individual channels within relatively short dwell times on the order of seconds. The possibility for partial disassembly of clusters stands in contrast to previous mathematical modeling of LTCC clustering (*14*), however a stochastic assembly process may hold true.

Interestingly, Ca_V_1.3 cluster positions in hiPSC-aCM were nearly immobile over long time scales. This is likely due to scaffolding at defined membrane locations, given the highly organized nature of cardiomyocytes including multi-protein CRUs with membrane tethering proteins like Junctophilin-2. In this line we confirmed that Ca_V_1.3, RyR2 and JPH2 consistently associate at the cell surface (Fig. 4), forming stable calcium release units, equivalent to peripheral dyadic junctions containing Ca_V_1.2 in ventricular cardiomyocytes (*57*). A direct interaction site between LTCC and JPH2 was recently postulated (*36, 58*), which is supported by our data showing strong colocalization of Ca_V_1.3 and JPH2 not only in hiPSC-aCM but also upon co-expression in HEK293 cells lacking cardiotypical proteins. In parallel these interactions could account for the observed fraction of immobile Ca_v_1.3 channels in SPT. However, our Cholesterol and Cav3 stainings did not reproduce a CRU association with lipid rafts (*59*).

Interestingly, previous studies reported the association of potassium channels K_V_2.1, K_Ca_1.1 and K_Ca_3.1 with neuronal CRUs, which further promoted LTCC clustering and function through direct interactions (*8, 60, 61*). This demonstrates that CRUs are dynamic, heterogenous structures enabling multifold protein interactions, in line with the emerging clustering model for LTCC. As indicated additionally by significant channel spacing and mobility, clustered Ca_V_1.3 channels are presumably intermixed with relevant interactors in the same nanodomain. This organization enables efficient channel regulation through transient, rather than constitutive interactions. Whether correlations exist between spatial arrangements and dynamics for distinct CRU constituents remains speculative but appears to be likely. Analogously these principles have been more extensively researched for Ca_V_2 channels in neuronal systems, however underlying a different functional context (*62, 63*). Notably, the confinement of presynaptic Ca_V_2 channels to cluster domains was shown to be dependent on alternative C-terminal splicing (*64, 65*) and similar organizational mechanisms were found for K_V_ channel clustering (*66, 67*).

For Ca_V_1.3 channels, there are two broadly expressed splice variants: Full-length, canonical Ca_V_1.3_42_ and C-terminally truncated Ca_V_1.3_42A_, which have distinct electrophysiological properties and a tissue-specific relative abundance, implying a cell-context specific fine-tuning of channel activation (*68*). Interestingly, endogenous cytosolic peptides of Ca_V_1.3 distal C-termini (DCT) competitively bind to Ca_V_1.3 channels and downregulate their function (*69*). To assess the relevance of Ca_V_1.3 C-termini for cluster formation, we took advantage of our flexible hiPSC-aCM expression system and introduced novel, artificial constructs of membrane-anchored Ca_V_1.3 C-terminal tail (CTT) for live-cell imaging. Cluster analysis surprisingly revealed that Ca_V_1.3 CTT intrinsically formed cell-surface clusters. Moreover, cluster abundance was clearly reduced for the shorter CTT sequence (180 aa of Ca_V_1.3_42A_) as opposed to the canonical CTT sequence (694 aa of Ca_V_1.3_42_) (Fig. 5). This clearly demonstrates that while the proximal, structured C-terminus is sufficient for cluster formation, there are additional protein interactions on the extended CTT of Ca_V_1.3_42_ that appear to enhance cluster formation or maintenance in the plasma membrane. Given that we observed a similar experimental outcome for ectopic expression in HEK293 cells (Fig. S7), it seems that the required protein interactions underlying this effect are not specific to cardiomyocytes.

This leads us to a comparison of putative protein interactions on the proximal versus extended (fulllength) CTT to relate our overall findings to possible clustering mechanisms. The extended CTT of Ca_V_1.3_42_ contains binding sites for JPH2, AKAP family and PDZ-binding proteins, which are relevant scaffolds for LTCC and may spatially define cluster nanodomains or mediate channel tethering (*5, 7, 8, 70*). While these interactions may enhance clustering, none of them appear to be strictly required for clustering, given that we showed cluster formation for the short CTT sequence of Ca_V_1.3_42A_. This proximal, structured C-terminus only contains the EF-Hand and IQ domains, which are important binding sites for modulation by CaM and CaBP (*71*). CaM binding was previously reported to mediate Cterminal channel coupling through dimer-like bridging (*4*) and confirmed to be transient in nature (*68*). Whether this mechanism is sufficient for multifold channel associations beyond dimerization was so far unknown but is supported by our data showing clustering of short Ca_V_1.3 CTT constructs. However, we observed a strongly (∼ 50%) reduced clustering of the short versus long CTT construct. This indicates that a mechanism involving CaM may be sufficient to form clusters, but not stabilize them, leading to disassembly or internalization. This agrees with an earlier coupled gating Ca_V_1.2 model that a mechanism involving CaM may be sufficient for cluster formation, but additional protein interactions on the extended distal C-terminus may increase cluster density through unknown mechanisms, possibly by increasing cluster stability. Interestingly, in tsA-201 cells and primary neurons (*4, 5*), the equivalent channel isoform Ca_V_1.3_42A_ showed only a mild (∼ 20%) reduction of cluster areas compared to full-length Ca_V_1.3_42_. The small effect size compared to our study of corresponding CTT constructs indicates that additional, CTT-independent mechanisms may increase the stability of Ca_V_1.3 channel clusters. Further investigation of clustering mechanisms and their functional impact using super-resolution and live-cell imaging studies presents a promising future direction for the research of LTCC regulation and pathophysiology.

In conclusion, we established a novel approach for researching LTCC clustering based on N-terminal channel tagging in hiPSC-aCM tailored for innovative multiscale super-resolution microscopy. Based on complementary results including DNA-PAINT, live-cell STED and SPT data, we propose that Ca_V_1.3 channel clusters consist of relatively mobile individual channels with large interspacing in defined membrane domains, which facilitate transient channel interactions with regulatory and scaffolding proteins to effectively regulate calcium signaling.

## Materials and Methods

### Plasmids

Ca_V_1.3 human cDNA (accession number NM_001128840.2, UniProt Q01668-1) was de-novo synthesized including an N-terminal ‘GGS’ linker. This cDNA was assembled into a vector encoding Nterminal fusion to the HaloTag (pHTN_HaloTag_CMV-neo, Promega G7721) using restriction cloning, yielding the Halo-Ca_V_1.3 plasmid. To alternatively generate an N-terminal mEGFP fusion, HaloTag was exchanged by restriction cloning to yield the GFP-Ca_V_1.3 plasmid.

A sequence encoding N-terminal HaloTag fused to the transmembrane sequence of non-clustering Integrin β_1_ (NM_002211.4) was generated as described by Svendsen, Zimprich, McDougall, Klaubert and Los (*38*) and assembled into an EFS-promotor driven vector (pRP-EFS, Vectorbuilder) to generate the control construct ‘HaloTM_Ctrl’. For Ca_V_1.3 CTT constructs, the insert sequence was fused with cDNA encoding CTT sequence of either human Ca_V_1.3_42_ (NM_001128840.2) or Ca_V_1.3_42A_ (XM_047448874.1), starting at amino acid position D1468, generating the long and short CTT plasmids ‘HaloTM_Cav13CT-L’ and ‘HaloTM_Cav13CT-S’.

### Cell culture and transfection

Human induced pluripotent stem cells (hiPSC) that were derived from healthy human donor cells (UMGi014-C clone 14 ‘isWT1.14’, kindly provided by the UMG Stem Cell Unit) were cultured in StemFlex medium (Gibco A3349401) and differentiated to atrial cardiomyocytes according to an established protocol (*72*). Cardiomyocytes were purified in glucose-free selection medium after differentiation and then cultured in RPMI 1640 medium (Gibco 72400021) with B-27 supplement (Thermo Fisher 17504044) lacking antibiotics. All differentiations showed spontaneous contractility. For imaging experiments, cells were seeded on Matrigel-coated glass-bottom imaging dishes (ibidi 81158) at subconfluent density. Two days after seeding, growth medium containing 2 μM CHIR99021 (Merck 361559) and 10% fetal bovine serum (Gibco 16140071) were added to enhance transfection efficiency (*41*). Cells were transfected the following day using 0.6–1 μg of plasmid and Viafect reagent (Promega E4981, 6 μl/μg plasmid). The medium was exchanged the following day to regular culture medium. Cells were imaged at 4–7 days after transfection.

HEK293 cells with constitutive expression of Ca_V_ subunits β_3_ and α_2_δ_1_ and inducible expression of α_1D_ (Charles River Laboratories CT6232) were cultured in DMEM/F12 medium containing selection antibiotics and 0.6 μM isradipine (Sigma I6658). For imaging experiments, cells were seeded on fibronectin-coated glass-bottom imaging dishes (ibidi 81158) in a growth medium lacking selection antibiotics. Transfection was performed using Lipofectamine 3000 reagent (Invitrogen L3000008) according to the manufacturer’s instructions with 0.6–1 μg plasmid DNA per imaging dish. A washing step was performed 3 h after transfection using a fresh culture medium. Microscopy experiments were carried out 1–2 days after transfection. For electrophysiology, the same protocol was applied to 6-well plates with 2 μg plasmid being added per well. The induction of α_1D_ subunit expression with Tetracycline was generally not performed, unless indicated.

### Cell labeling for microscopy

For Halo-Ca_V_1.3 imaging by confocal and STED microscopy, a labeling solution containing 100 nM JF646-HTL (Promega GA1121) in phenol red-free culture medium was freshly prepared. Live-cell labeling was performed by incubation of hiPSC-aCM in labeling solution for 30 min at 37°C, optionally followed by co-labeling with Cholesterol-Star488/ -StarOrange (Abberior 0206, 40 nM) for 10 min or ER-Tracker Red (Invitrogen E34250, 1 μM) for 30 min in cell culture medium at 37°C. After labeling, a wash-out step was performed by incubation with fresh culture medium for 2 h at 37°C. Afterwards, cells were washed thrice and imaged in live-cell imaging solution (Thermo Fisher A14291DJ).

Dual channel labeling for SPT experiments was performed in phenol red-free culture medium: First, a solution of 250 pM JF646-HTL was applied for 10 min at 37°C. Then, a solution containing 50 nM JF549-HTL (Promega GA1110) was applied for 15 min at 37°C. This was followed by two washing steps and incubation in culture medium for 2 h at 37°C to achieve effective wash-out of unbound ligands. The cells were then washed four times with live-cell imaging solution for 5 min each and subsequently imaged.

For confocal imaging of CTT constructs, hiPSC-aCM or HEK293 cells were incubated with a labeling solution containing both cell-impermeant HTL-Alexa488 (Promega G1001, 1 μM) and Cellmask Deep Red Plasma Membrane stain (Invitrogen C10046, 5 μg/mL) in phenol-red free culture medium for 10 min at 37°C. Then, cells were washed twice with medium and twice with live-cell imaging solution before imaging. Cellmask stain was optionally exchanged for Cholesterol-PEG-KK114 (*37*), yielding similar membrane staining. Notably, alternative HaloTag labeling using Alexa660-HTL (Promega G8471) at 3.5 μM concentration did not generate a sufficient labeling outcome.

For immunofluorescence of cells expressing Halo-Ca_V_1.3, live-cell incubation with JF646-HTL was performed as described above. Afterward, cells were washed twice with phosphate buffered saline (PBS) and fixed for 10 min by 4% paraformaldehyde (PFA) diluted in Dulbecco’s PBS containing Ca^2+^ and Mg^2+^ (DPBS; Gibco 14040083), then blocked and permeabilized for 1 h in blocking buffer (10% bovine calf serum and 0.1% Triton X-100 in DPBS). Cells were then incubated with primary antibody diluted in blocking buffer overnight at 4°C. This was followed by a secondary antibody incubation in a blocking buffer for 90 min at room temperature. Cells were imaged in DPBS, SlowFade Diamond (Invitrogen S36967) or ProLong Gold (Invitrogen P36930).

DNA-PAINT labeling of GFP-Ca_V_1.3 transfected cells was carried out according to the immunofluorescence protocol, but cells were fixed in 4% PFA for 20 min at RT, both before and after the labeling procedure to achieve post-fixation. The blocking buffer was supplemented with 0.1 mg/mL sheared salmon sperm DNA (Thermo Fisher 15632011) and 0.05% w/v dextran sulfate (Merck D4911) and Image-iT FX reagent (Invitrogen I36933) was applied for 10 min after blocking to reduce nonspecific imager binding (*73, 74*). GFP nanobody conjugated to R4-docker DNA (sequence 5’ → 3’: ACACACACACACACACACA, Metabion) was applied in dilution buffer (3% BCS, 0.1% Triton X-100, 0.05 mg/mL sheared salmon sperm DNA in DPBS) for 1 h at RT.

The following antibodies were used in this study:

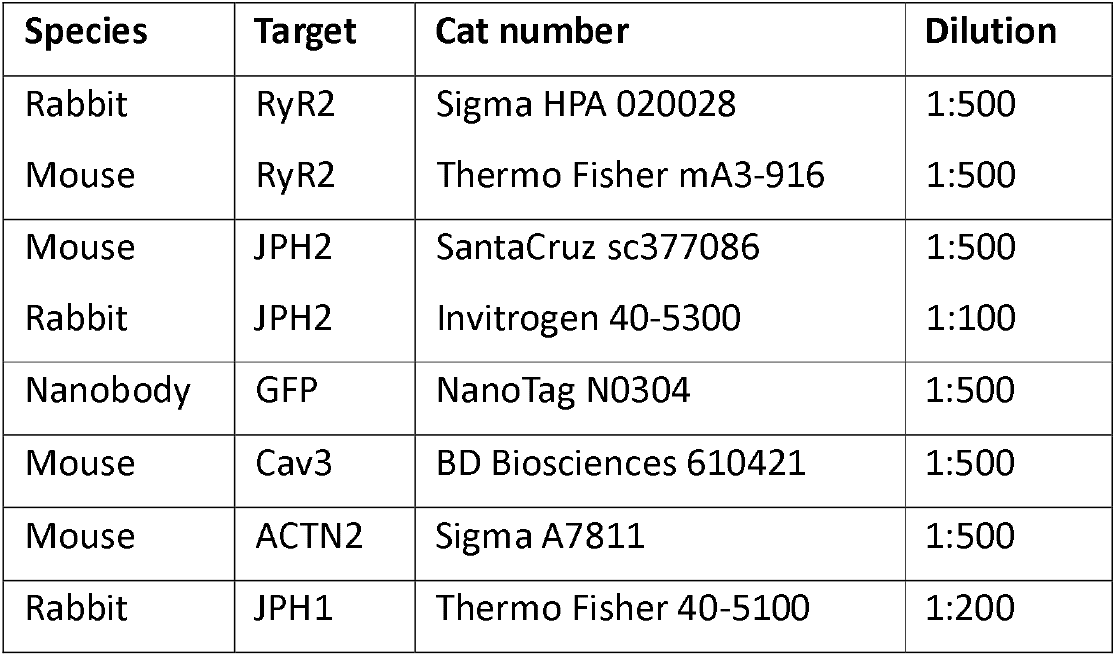

### Microscopy setup and image acquisition

Confocal and STED imaging were performed using an Abberior Expert Line inverted microscope equipped with an oil immersion objective lens (Olympus UPlanSApo 100x NA 1.4), pulsed excitation lasers at wavelengths 640/591/485 nm, pulsed STED laser at wavelength 775 nm, Abberior QUAD scanner and avalanche photodiode detectors (Excelitas Technologies SPCM-AQRH). Acquisition settings for quantitative JF646 STED imaging were as follows: 30% excitation laser power at 640 nm, 12% STED laser power, pixel size 25 nm, pixel dwell time 64 μs, time gating window 0.5–6 ns. Confocal images were generated at variable excitation powers depending on the experiment and a pixel size of 80 nm. Image channels were recorded separately in line steps to avoid fluorescence crosstalk. All imaging was performed at RT.

DNA-PAINT and SPT measurements were performed on a custom-built TIRF optical setup, as described elsewhere (*20*). The optical configuration is shown in Fig. S8. Briefly, 488 nm (Omicron PhoxX+ 488-100), 561 nm (Changchun MGL-FN-561-100), and 638 nm (Omicron PhoxX+ 638-150) lasers were used for sample excitation. A neutral density filter (Thorlabs NE10A-A) in tandem with the variable neutral density filter ND (Thorlabs NDC-50C-4-A) were used to adjust the laser excitation power. The laser beam was coupled into a single-mode optical fiber (SMF, Thorlabs P1-460B-FC-2) with a typical coupling efficiency of 40%. After exiting the optical fiber, the collimated laser beam was expanded by a factor of 3.6X using telescope lenses (TL1 and TL2). The typical excitation intensity at the sample was ∼ 1 kW/cm^2^ for high-photon flux DNA-PAINT imaging.

The laser beam was focused onto the back focal plane of the TIRF objective (Olympus UAPON 100X oil, 1.49 NA) using achromatic lens L1 (Thorlabs AC508-180-AB). Mechanical shifting of the beam with respect to the optical axis was done through a translation stage (TS, Thorlabs LNR25/M) to allow for a change between different illumination schemes: EPI, HILO, and TIRF. The smooth lateral positioning of a sample was achieved by using a high-performance two-axis linear stage (Newport M-406). In addition, an independent one-dimensional translation stage (Thorlabs LNR25/M) together with a differential micrometer screw (Thorlabs DRV3) was used to shift the objective along the optical axis for focusing on different sample planes. The spectral separation of the collected fluorescence light from the reflected excitation light was achieved using a multi-band dichroic mirror (DM, Semrock Di03 R405/488/532/635), which directed the emitted fluorescence light towards the tube lens L2 (Thorlabs AC254-200-A-ML). The field of view was physically limited in the emission path by an adjustable slit aperture (OWIS SP60) positioned in the image plane. Lenses L3 (Thorlabs AC254-100-A) and L4 (Thorlabs AC508-150-A-ML) were used to re-image the emitted fluorescence light form the slit onto an emCCD camera (Andor iXon Ultra 897). A band-pass filter (BPF, BrightLine HC 692/40) was used to further block the scattered excitation light. The total magnification of the optical system on the emCCD camera was 166.6X, resulting in an effective pixel size in the sample space of 103.5 nm.

DNA-PAINT acquisition was performed in a TIRF mode, with the exposure time of 30 ms and EM gain of 500. First, cells with GFP expression level were selected and then DNA PAINT movies of 30–50k frames were acquired. The imager concentration was in the range of 0.5–1 nM. Single particle tracking was performed in a TIRF mode with the exposure time of 30 ms and EM gain of 500 (JF646) or 100 (JF549). Typically, the acquisition of a single movie took 2–5 minutes. All experiments were done at 22°C temperature, which was crucial for the mechanical stability of the optical setup.

### Confocal and STED image analysis

Image analysis was performed in ImageJ Fiji (*75*) version 1.54f. All analyses were applied selectively to annotated cell areas. For STED image segmentation, the FFT bandpass filter (2.5–20 px) was applied to remove high-frequency noise and unstructured background signal. Candidate signal spots were identified in the filtered image by maxima detection and peak expansion to half-maximal intensity (FWHM) using the ImageJ plugin Interactive H-Watershed. Resulting candidate regions of interest (ROI) were discarded if containing less than 5 pixels or a mean intensity less than 50% above the background signal. All remaining ROIs representing specific signals were used for area and brightness measurements. Cluster diameters were calculated from segmented areas using 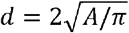 assuming circular shape. Signal brightness was measured in raw image data and corrected for local background by subtraction of the mean brightness of a ring-like ROI obtained by differential enlargement of each cluster ROI by 4 versus 2 pixels.

For molecular counting in STED images (Fig. S2), DNA Origami reference structures containing 23 ± 3 and 7 ± 1 JF646 dye binding sites were purchased from GATTAquant and immobilized on the surface of ibidi glass-bottom imaging dishes coated with BSA-biotin. A calibration measurement was performed by applying the same quantitative nanoscopy workflow as for Halo-Ca_V_1.3 cluster samples. The resulting distribution of single particle brightness was used to determine a single dye brightness value for molecular counting of channel molecules in cluster ROIs.

For confocal-based colocalization analysis, three-channel images were binarized using FFT bandpass filtering (4–40 px) followed by automated local thresholding (‘Otsu’, radius 20 px) in each channel. Colocalization was determined in raw image data as the fraction of above-threshold Ca_V_1.3 signal mass in binarized JPH2-, or RyR2-positive area, or JPH2-RyR2 double-positive area (as defined by Manders M1 colocalization coefficient). To analyze CRU composition, CRU signals were defined as the pixel-wise product of JPH2 and RyR2 signals. Then, the fraction of above-threshold CRU signal mass colocalized with binarized Ca_V_1.3 signals was calculated (corresponding to Manders M2).

For confocal-based cluster analysis of CTT constructs, images were smoothed by a Gaussian filter (σ = 1 px) and then binarized using automated thresholding (‘Moments’). Resulting spots were quantified by ‘Analyze Particles’ with particle size 6–120 px^2^. Larger particles were not analyzed as they were atypical for clusters and may originate from endosomes.

The ImageJ macro code used in this section is provided as Supplementary Software. DNA-PAINT image reconstruction and analysis

Raw DNA-PAINT image sequences were processed in ImageJ using the ThunderSTORM plugin (*76*). The following parameters were used for emitter localization:

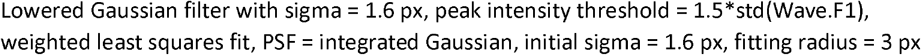

For drift and vibration correction, the recently published DME algorithm (22, version 1.2.1) was implemented using MATLAB R2022A. We increased the robustness of long-term drift tracking by applying DME iteratively for decreasing time bins, which performed better compared to redundant cross correlation (RCC), which was suggested in the original implementation. Our optimization efforts resulted in the following parameters:

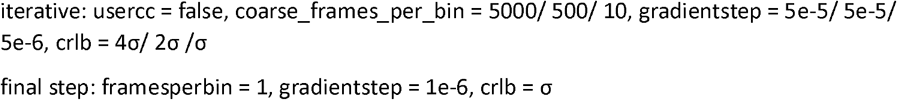

The obtained improvements in localization precision were evaluated by NeNa, which was applied to the central image region (code adapted from *23*). To further increase localization precision, detections were merged in ThunderSTORM with parameters *max. dist. = 30 nm, max. frames = 4, max. off frames = 0* and then filtered by the criteria *detections > 2 & uncertainty < 10 nm*. Subsequently, a density filter with *radius = 12 nm, n_min = 4* was applied to discard spurious signals. The reconstruction was then generated by Gaussian rendering with *sigma = 5 nm, magnification = 26*, resulting in 4 nm image pixel size.

For molecular mapping and cluster analysis, a custom-written analysis was applied using MATLAB R2022A. The analysis was limited to manually annotated cell footprints. Briefly, reconstructions were smoothed by the H-maxima transform (*imhmax*) followed by local maxima detection

(*imregionalmax*), thus detecting putative channel positions by combining localizations and binding events originating from the same binding site. These positions were then subjected to DBSCAN clustering with *minPts = 3 and ε = 100 nm (see Fig. S3C)*. The points comprising each cluster were then outlined using the *boundary* function. The resulting polygons were expanded by *d = 10 nm* using the *polybuffer* function to account for physical channel dimensions and localization error. Nearestneighbor distances across all points or only clustered points were determined using the *knnsearch* function.

The MATLAB code used in this section is provided as Supplementary Software

### Single particle tracking and motion analysis

Localization and tracking of single-molecule image series was performed using the *TrackIt* package (*56*) running in MATLAB R2022A. The following parameters were used:

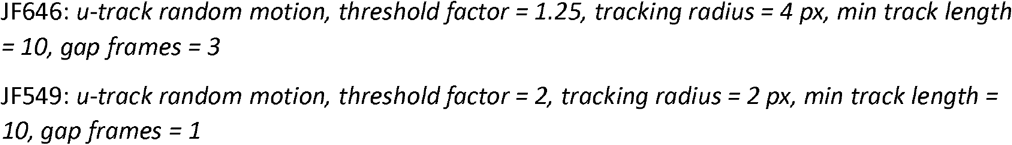

Mean jump distance (MJD) and diffusion coefficient (D) were determined for each track using the same software. D values were derived using a linear fit of the first 5 MSD values based on the equation *MSD*(*τ*) = 4 *D τ* + *b* where MSD is the mean squared displacement over all localization pairs for any given lag time *τ* (based on the frame interval), and b is an offset term reflecting the experimental localization error (*27*). For confocal time series in Fig. S4 containing relatively long tracks, the logarithmic fit function *MSD*(*τ*) = 4 *D τ* ^α^+ *b*. was applied, where α is an exponent indicative of the motion type (*77*). For visualization, tracks were rendered with temporal time coding using the color map *turbo*.

Confinement metrics were calculated for each track containing at least 10 localizations by averaging over a sliding time windows of 300 ms duration. Convex hull areas were calculated using the *convhull* function. The radius of gyration (R_Gyr_) was calculated as the root mean square of pointwise distances to the center of mass. Notably, an apparent motion of immobile particles is always detected in experimental SPT data due to localization errors. Mathematical modeling of this effect was used to estimate the influence on our measurements. We determined an approximate localization precision (σ) of 25 nm in our SPT images using ThunderSTORM. Subsequently, the following formulae (*33*) were used to determine MJD and R_Gyr_ values for immobile particles:

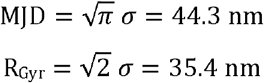

To generate value distributions resembling our experiment as a point of comparison, we performed a Monte-Carlo simulation (n = 10,000 runs) of immobile particles according to the experimental track length distribution (Fig. S5B) and estimated localization precision σ = 25 ± 5 nm, resulting in:

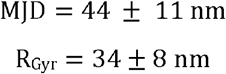

The MATLAB code used in this section is provided as Supplementary Software.

### Patch clamp measurements

HEK293 CT6232 cells were seeded on fibronectin-coated 6-well plates and transfected with 2 μg of plasmid per well encoding Halo- or GFP-tagged Ca_V_1.3 as described earlier. For control conditions, Ca _V_1.3^WT^ (α_1D_) was induced by adding 1 μg/mL Tetracycline (Sigma T7508) to the culture medium in parallel to transfections. Cells were harvested one day after transfection as follows: After washing with pre-warmed PBS, cells were dissociated by incubation with 0.25% Trypsin-EDTA solution (Thermo Fisher 25200056) for 30 s at 37°C. The cells were then suspended in growth medium and centrifuged at 100 g, 4°C for 5 min. The cell pellet was then resuspended in divalent-free HBSS (Gibco) at 4°C. Automated patch-clamp experiments were conducted with the SyncroPatch 384 (Nanion Technologies GmbH) device with thin borosilicate glass, single aperture 384-well chips (NPC384T 1 x S-type). Application of negative pressure (150–250 mbar) allowed for whole-cell access. To measure IV curves, Ca_V_1.3 currents were elicited with a voltage-step protocol at 0.5 Hz with a holding potential of -80 mV followed by a 100 ms test-pulse at -40 mV to +70 mV with 5 mV increments. A 100 ms depolarizing ramp from -80 mV to -40 mV has been used as pre-pulse in experiments, in which Ca^2+^-current kinetics were quantified. Experiments were carried out at 22–24 °C. Internal solution contained (in mmol/L): EGTA 10, HEPES 10, CsCl 10, NaCl 10, CsF 110, pH 7.2 (with CsOH). Bath solution contained (in mmol/L): HEPES 10, NaCl 140, glucose 5, KCl 4, CaCl_2_ 2, MgCl_2_ 1, pH 7.4 (with KOH). Currents were recorded with a Tecella amplifier controlled by PatchControl 384 software. Recordings were analyzed offline with DataControl 384 software (both: Nanion Technologies GmbH) (*78*).

### General data analysis

Data was visualized and statistically analyzed using GraphPad Prism version 10 and MATLAB R2022A.

## Supporting information

Supplementary Software

## Acknowledgements

We are grateful for the excellent technical assistance by Timo Schulte, Brigitte Korff and Elli Rehbein-Bode. Stephan E. Lehnart is a principal investigator of DZHK (German Centre for Cardiovascular Research).

## Author contributions

Conceptualization: NS, TK, JE, SEL

Methodology: NS, RT, TK, FS, IS

Software: NS

Investigation: NS, RT, SB, FS, RK, IS

Formal analysis: NS, RT

Visualization: NS, SB

Writing – original draft: NS

Writing – review & editing: NS, TK, SEL

Supervision: TK, NV, JE, SEL

Funding acquisition: SEL, JE, NV

## Data and materials availability statement

All data needed to evaluate the conclusions in the paper are present in the paper and/or the Supplementary Materials.

The generated code for image data processing and analysis is provided as Supplementary Software.

