## Supplementary Figures for "Nanoscale architecture and dynamics of Ca_V_1.3 channel clusters in cardiac myocytes revealed by single channel nanoscopy"

Software files correspond to experiments of the main text figures as follows:

Fig. 1: „ImageJ_1 STED CaV cluster analysis.ijm“.
Fig. 2: „DNA_PAINT_1_DME_drift_correction.m“;
„DNA_PAINT_2_MolecularMapping.m“;
„DNA_PAINT_3_ClusterAnalysis.m“.
Fig. 3: „SPT_1_Diffusion_analysis_trackit.m“;
„SPT_2_SimulationImmobileLocError“.
Fig. 4: „ImageJ_2 Confocal CRU colocalization.ijm“.
Fig. 5: „ImageJ_3 Confocal CTT cluster thresholding.ijm“.

Functionality is further described in the Methods section and the in-file comments.

**2) Supplementary Figures S1 – S8: see below**

**Figure S1: Automated patch clamp measurements confirm similar electrophysiological characteristics of wild-type and Halo- or GFP-tagged $Ca_V1.3$ channels expressed in HEK293 CT6232 cells.**

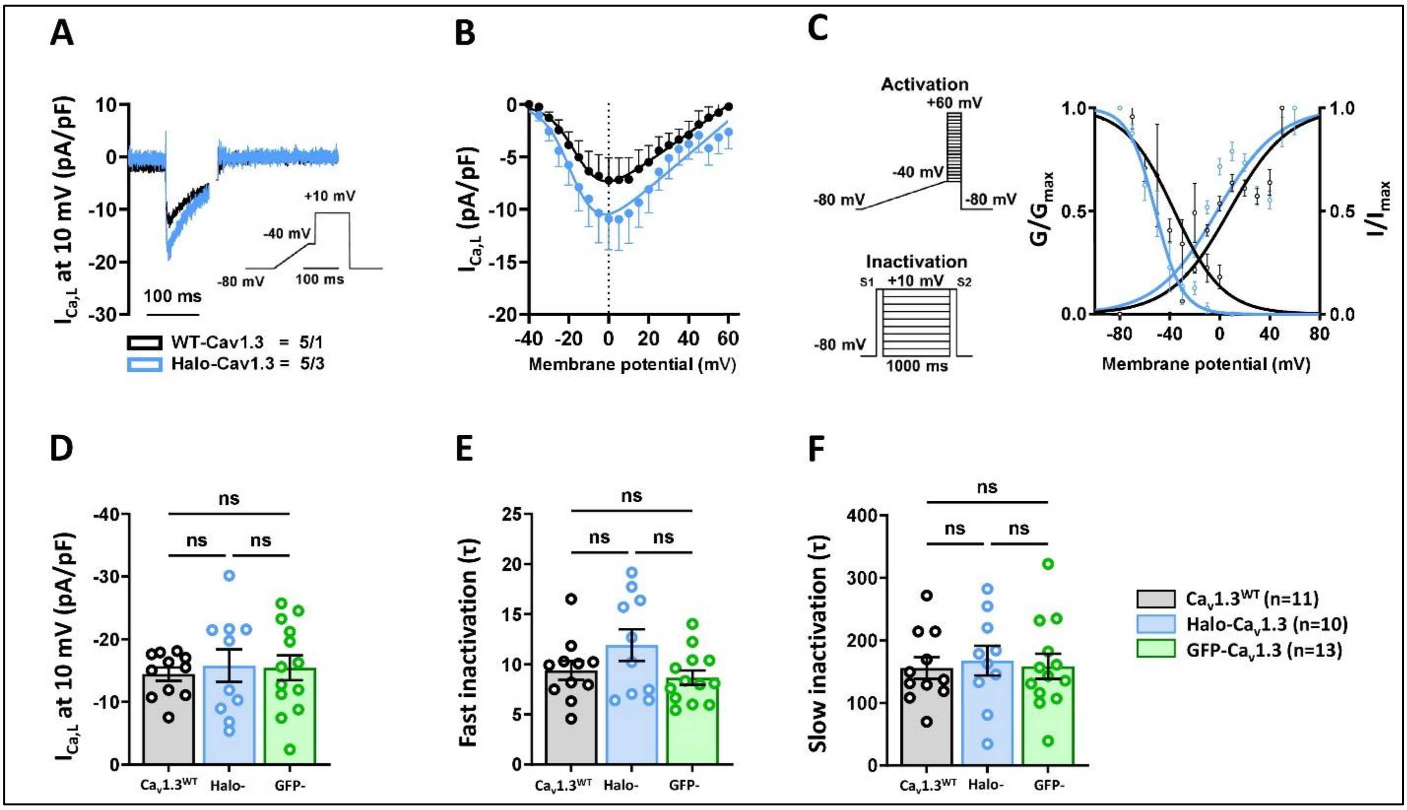


HEK293 CT6232 cells expressing the accessory $Ca_V$ channel subunits $\alpha_2\delta_1$ and $\beta_3$ were induced to express pore-forming, wild-type $\alpha_{1D}$ ($Ca_V1.3^{WT}$) or transfected with a plasmid encoding the tagged $\alpha_{1D}$ (Halo-$Ca_V1.3$). Whole-cell calcium currents were measured using the Nanion SyncroPatch 384 device.

In a first set of experiments Halo-tagged $Ca_V1.3$ channels were compared to WT channels **(A-C)**, showing similar electrophysiological characterisics: **A,** $Ca_V1.3^{WT}$ and Halo-$Ca_V1.3$ cells show typical $I_{Ca,L}$ membrane current ($I_M$) with the shown voltage-ramp protocol. **B,** Current–voltage (I–V) relationship curves for $I_{Ca,L}$ WT-$Ca_V1.3$ and Halo-$Ca_V1.3$ cells have similar shapes, while different amplitudes indicate different expression levels. **C,** $Ca_V1.3^{WT}$ and Halo-$Ca_V1.3$ cells showing similar current activation and inactivation. Note the voltage-ramp protocols for I–V/activation experiments and S1/S2 voltage protocols for $I_{Ca,L}$ inactivation. The graph shows similar $I_{Ca,L}$ activation ($G/G_{max}$) in $Ca_V1.3^{WT}$ and Halo-$Ca_V1.3$, with corresponding inactivation ($I/I_{max}$) curves. (n = number of WT-$Ca_V1.3$ cells, 5 from 1 batch, and Halo-$Ca_V1.3$ cells, 5 from 3 batches. The I–V curves were fitted with a modified Boltzmann equation.)

In a second set of experiments both GFP- and Halo-tagged $Ca_V1.3$ channels were compared to WT channels **(D-F)**, with similar electrophysiological characterisics: Peak $I_{Ca,L}$ amplitudes (pA/pF) (**D**) and biphasic inactivation kinetics of $I_{Ca,L}$ analyzed as fast inactivation (**E**) and slow inactivation (**F**) were not statistically different (ns). All parameters were compared using ANOVA Kruskal-Wallis test with Dunn's correction, n = number of cells, data indicate mean ± SEM.

**Figure S2: Brightness referencing method for molecular counting of JF646 fluorophores.**

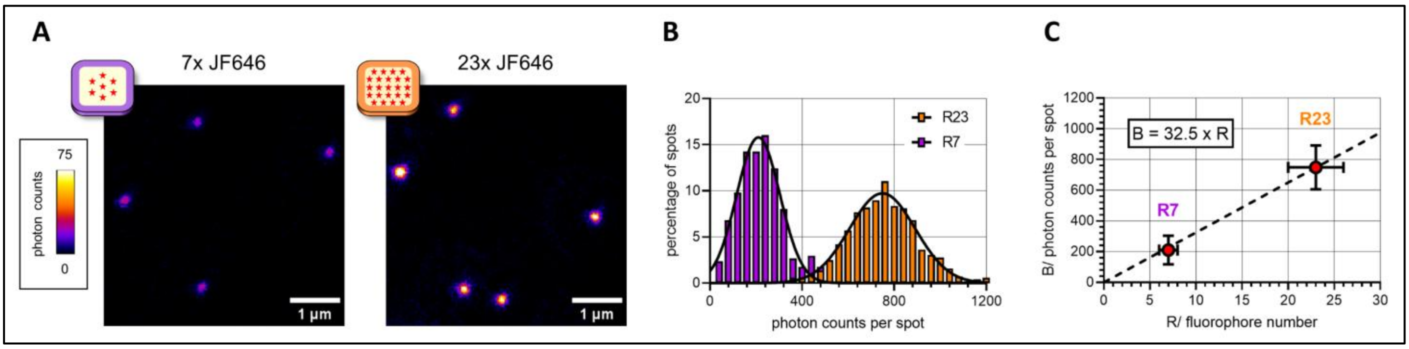


DNA Origami linked to 7 or 23 JF646 dye molecules were immobilized on coverslips and recorded by STED imaging under equal conditions as for cellular Halo-$Ca_V$1.3 cluster imaging **(A)**. By image analysis of spot-like signals, a distribution of integrated photon counts across all detected spots was determined for each sample **(B)**. The histograms were fitted by normal distributions to retrieve mean brightness values, which was used for a linear fit of spot brightness to dye molecule counts in **(C)**. The determined conversion factor (32.5 photon counts per fluorophore) was used for image analysis of Halo-$Ca_V$1.3 samples to retrieve labeled channel counts within clusters.

**Figure S3: Optimization of DNA-PAINT image reconstruction and DBSCAN clustering.**

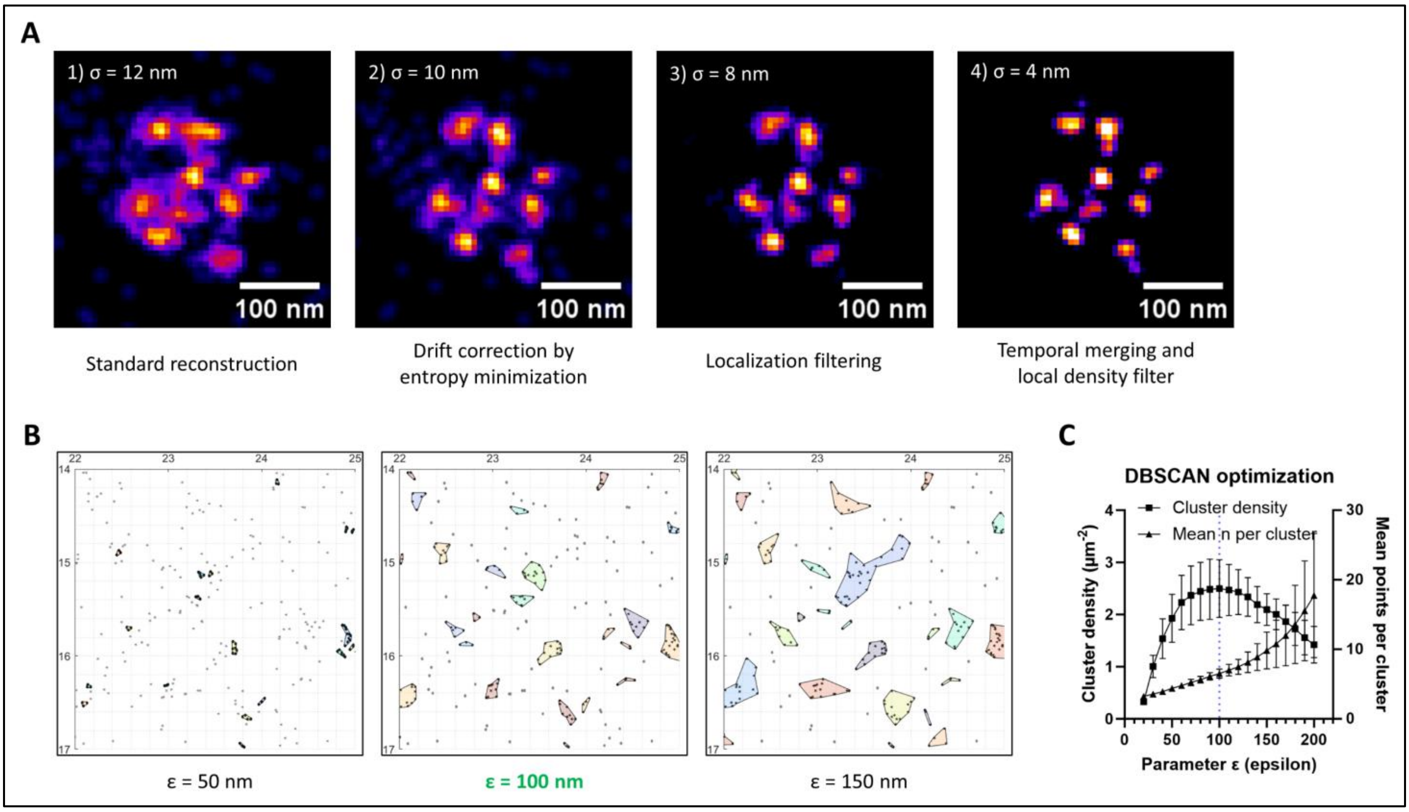


**A)** Representative reconstruction of a GFP-Ca$_V$1.3 channel cluster imaged by DNA-PAINT in TIRF mode. A successive improvement of the localization-based image reconstruction over standard reconstruction (1) was achieved by applying a customized version of drift correction by entropy minimization (DME, 2), followed by either histogram-based localization filtering (3), or followed by temporal merging of subsequent localizations and local density filtering (4).

**B)** The point clustering algorithm DBSCAN was applied to DNA-PAINT molecular map data. Three exemplary values for the parameter ε give rise to distinct clustering results.

**C)** Graph showing the change in DBSCAN cluster density and points per cluster as a function of ε parameter values at minPts = 3. For ε = 100 nm, the highest cluster density (2.5 µm$^{-2}$) is observed, while higher ε values lead to merging of pre-existing clusters and increased heterogeneity.

**Figure S4: Confocal timelapse imaging demonstrates immobility of Halo-$Ca_V$1.3 clusters across time scales.**

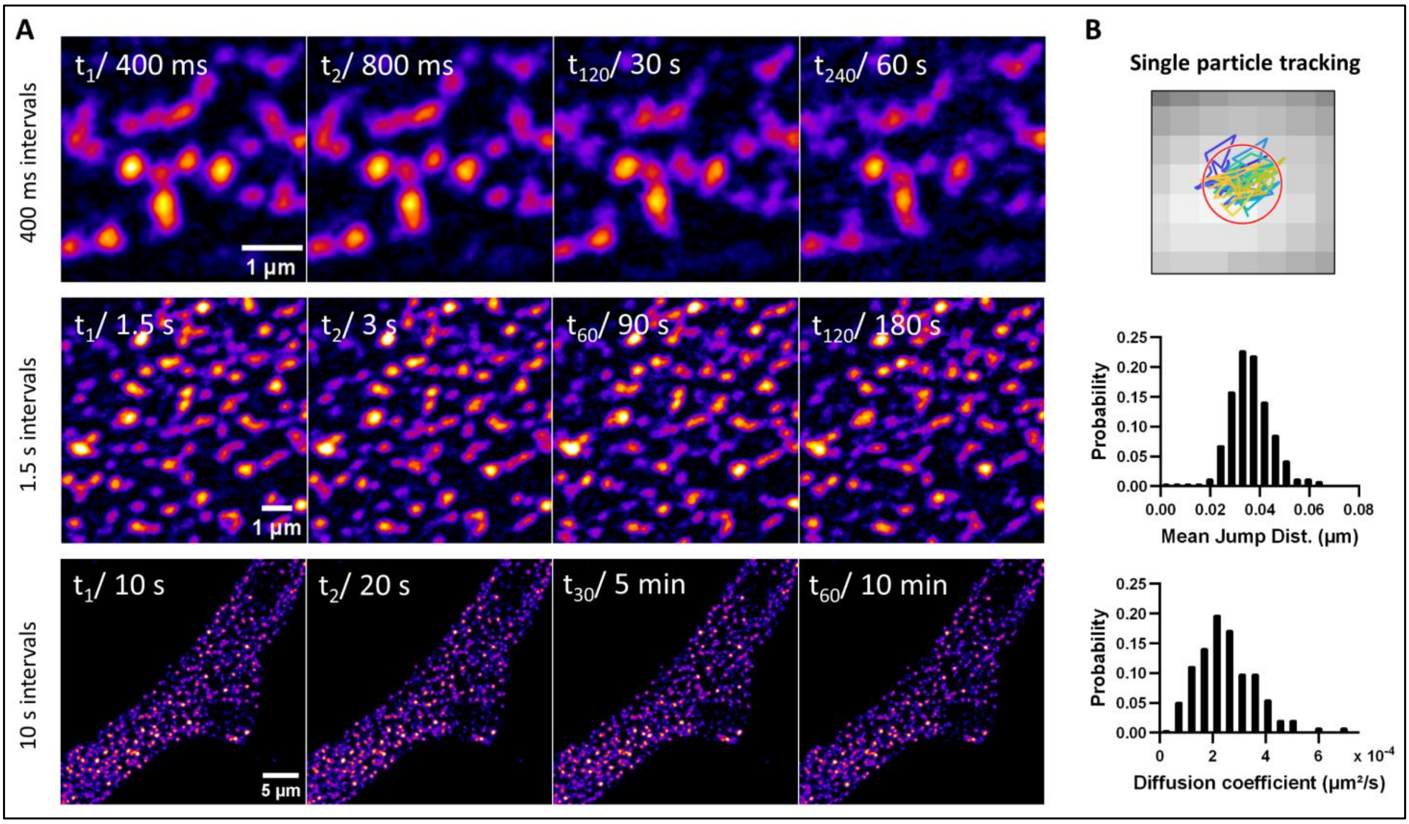


**A)** Confocal timelapse imaging of hiPSC-aCM expressing Halo-$Ca_V$1.3 (row 1+2) or GFP-$Ca_V$1.3 (row 3) shows cluster positions in the basal plasma membrane. Images series were recorded in intervals of 400 ms, 1.5 s and 10 s. For each timelapse, the first, second, middle and last frame are shown.

**B)** Representative trajectory of a single cluster position shown on a 30 nm pixel grid, generated by SPT of a timelapse at 1.5 s intervals and 30 nm pixel size. Quantitative analysis of SPT data reveals low jump distances of ~ 35 nm reflecting the localization uncertainty and MSD fit-derived diffusion coefficients of less than $10^{-4}$ µm²/s, thus confirming immobility of the tracked cluster positions.

**Figure S5: Supporting data for single particle tracking analysis.**

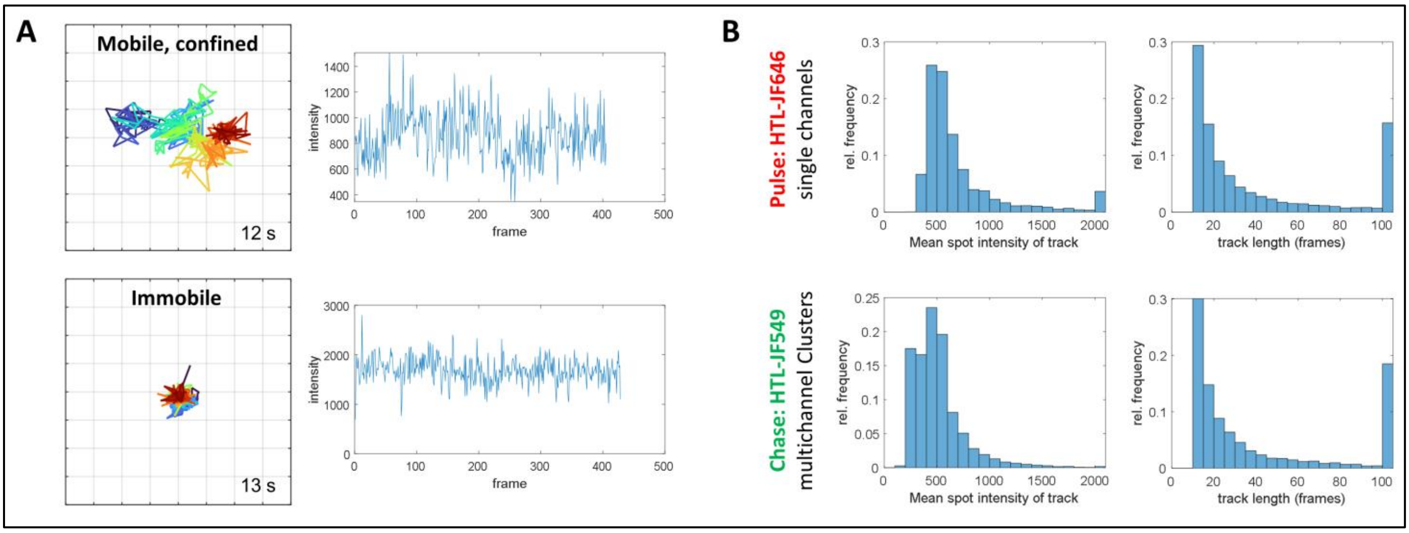


**A)** Intensity time traces for the exemplary tracked spots shown in Fig. 3C. No bleaching steps were observed in the majority of long tracks.

**B)** Mean spot intensity and track length distributions indicative of tracking performance were calculated. Both metrics show similar distributions for both imaging modes, which excludes a potential bias in the comparative diffusion analysis.

**Figure S6: Halo-Ca$_V$1.3 colocalization with nanodomain and compartment markers.**

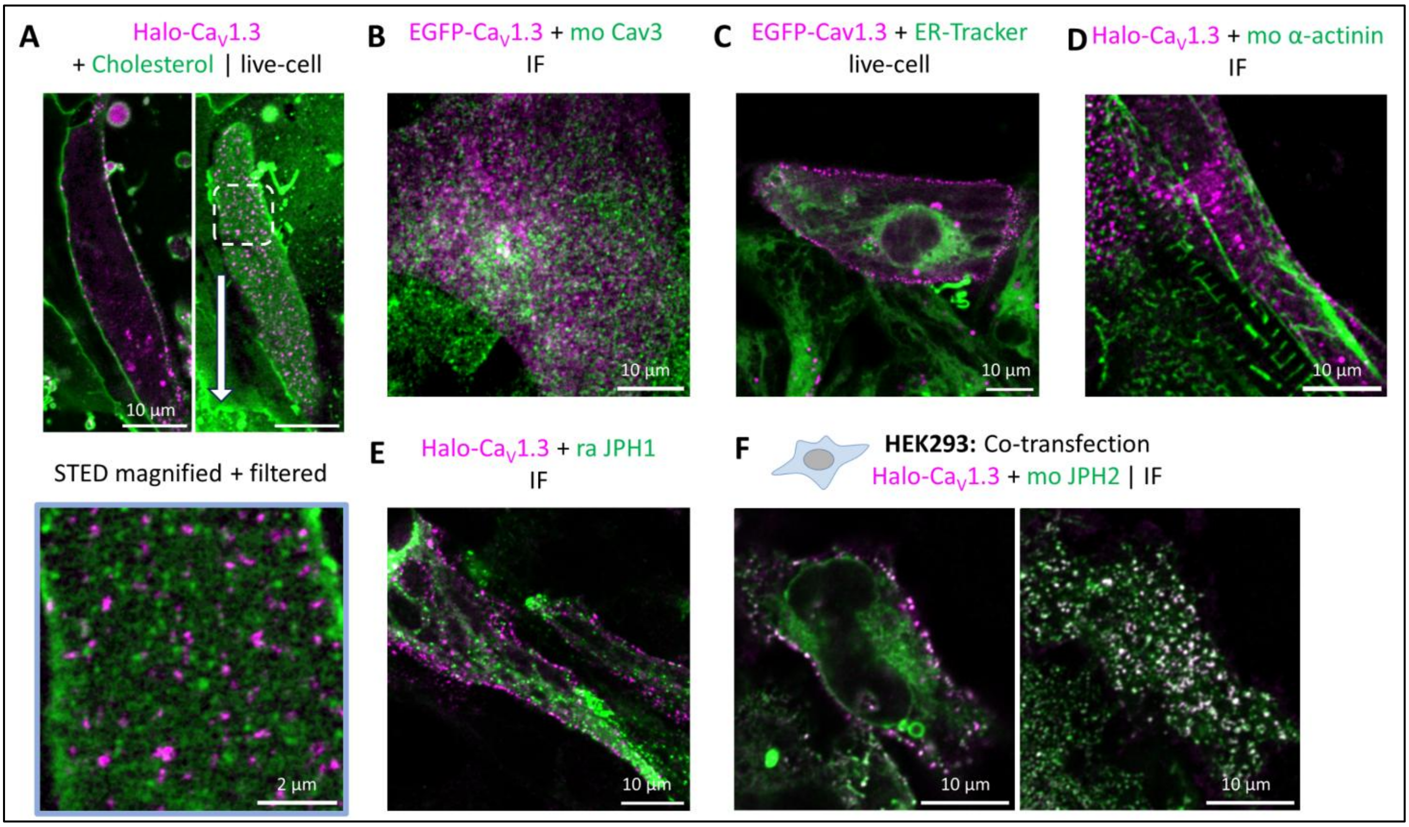


**A)** Live-cell confocal images of hiPSC-aCM expressing Halo-Ca$_V$1.3, labeled by HTL-JF646 and Cholesterol-StarOrange. Ca$_V$1.3 clusters and Cholesterol both localized to the plasma membrane (top), but dual-channel STED imaging in the basal membrane focal plane (bottom) showed rather exclusion-like arrangement with Cholesterol-containing nanodomains.

**B)** Immunofluorescence of hiPSC-aCM expressing EGFP-Ca$_V$1.3 showed rather low colocalization with Caveolin-3 (Cav3).

**C)** Live-cell imaging of hiPSC-aCM showed a mutually exclusive distribution of EGFP-Ca$_V$1.3 and endoplasmic reticulum, labeled by ER-Tracker Red.

**D)** Immunofluorescence of hiPSC-aCM expressing Halo-Ca$_V$1.3 showed no colocalization with cardiac α-actinin or Junctophilin-1 (JPH1) **(E)**.

**F)** Immunofluorescence of HEK293 CT6232 cells transfected with Halo-Ca$_V$1.3 and JPH2-CFP showed extensive colocalization of clustered spots in the basal membrane focal plane.

**Figure S7: $Ca_V1.3$ C-terminal construct expression in HEK293 leads to cluster formation independent of the cardiac proteome.**

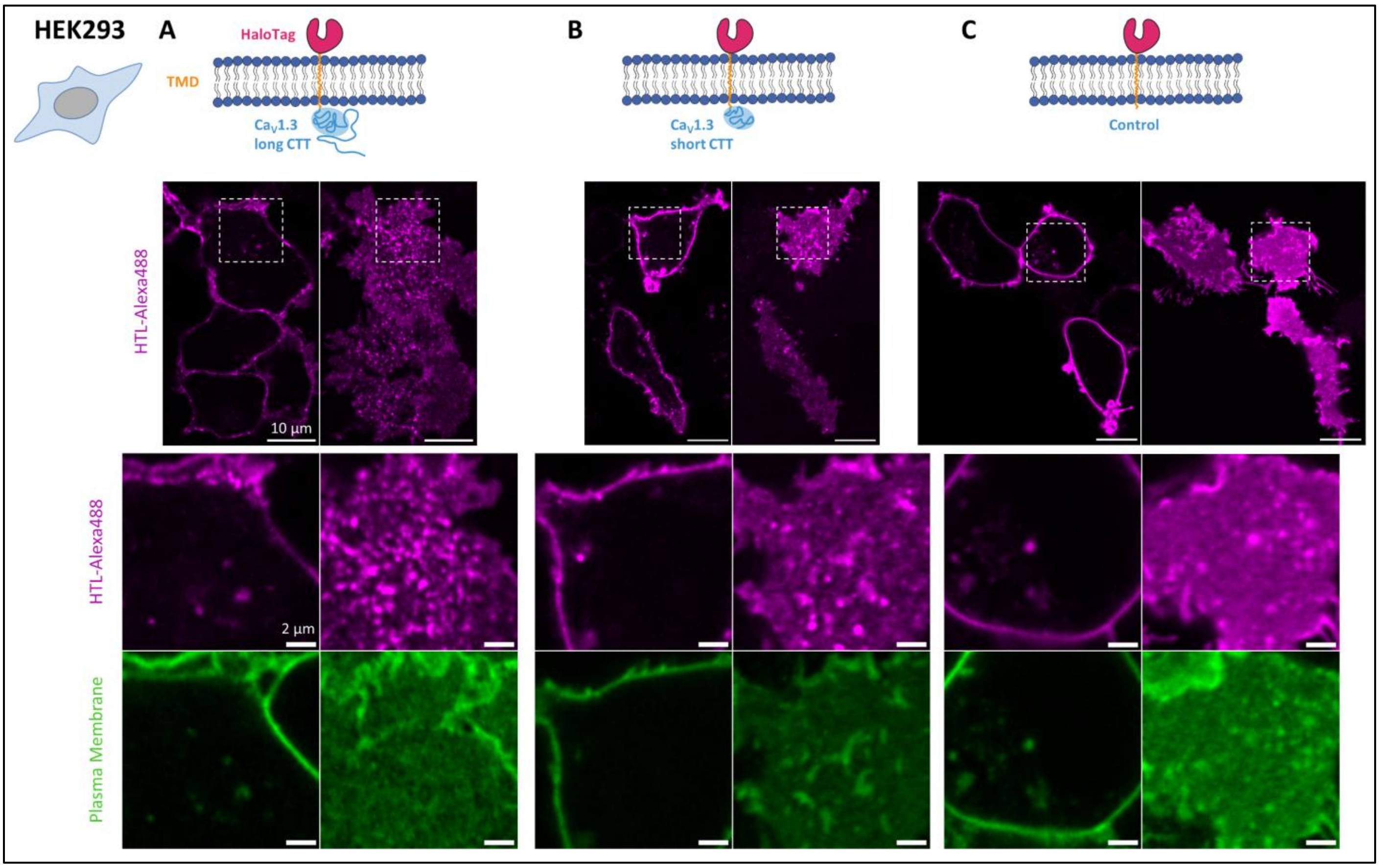


**A)** $Ca_V1.3$ C-terminal cytosolic tail (CTT, long isoform) fused to cell-surface HaloTag was expressed in HEK293 CT6232 cells and labeled with cell-impermeable HTL-Alexa488. The cells were co-stained with the plasma membrane marker Cholesterol-PEG-KK114 and imaged by live-cell confocal microscopy.

**B)** Expression of the equivalent fusion protein containing the short C-terminal tail splice variant.

**C)** Expression of a control construct containing only cell-surface HaloTag without CTT sequence, serving as a negative control.

**Figure S8: Illustration of the custom-built optical setup (described in the Methods section)**

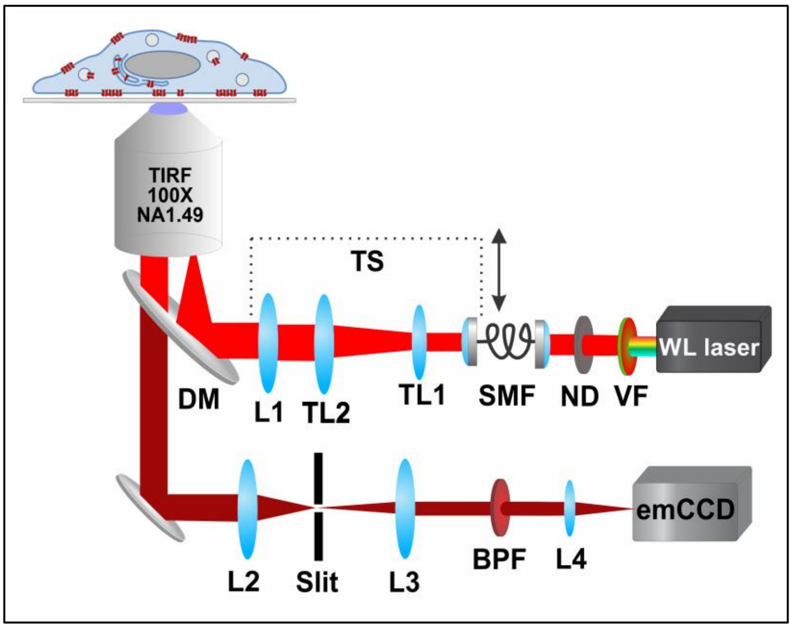
