## Supplementary Software for "Nanoscale architecture and dynamics of Ca_V_1.3 channel clusters in cardiac myocytes revealed by single channel nanoscopy": DNA_PAINT_1_DME_drift_correction.m

```
% Requires: DME package v1.2.1 (https://github.com/qnano/drift-estimation, Cnossen et al. 2021)
% Requires: File 'commonFunctions.m' for NeNa precision estimate (https://github.com/Endesfelder-Lab/SMLMComputational, Martens et al 2021)
% Input: ThunderSTORM localization table (name.csv) without DC or other processing
% Output: Drift corrected localization table, drift curves, uncertainty estimate, log file 

%% Setup and file processing
clearvars
warning('off', 'MATLAB:MKDIR:DirectoryExists');
warning('off', 'MATLAB:loadlibrary:cppoutput');
[file.name,file.folder] = uigetfile('*.csv','Select a file');
if contains(file.name, 'DME')
    warning('file %s seems to be already corrected', file.name);
end
fullpath = strcat(file.folder, '\', file.name);
[dir,name,ext] = fileparts(fullpath);
subfolder = '\DME\';
mkdir(strcat(dir, subfolder));
diaryfile = strcat(dir, subfolder, name, '.txt');
diary(diaryfile)

%% Read ThunderSTORM localization table
%  Cave: Last table header is named 'uncertainty_xy [nm]' in another Thunderstorm version (needs 3x renaming in script) 
%        and is missing in the 'phasor' version of ThunderSTORM
A0 = readtable(fullpath,"VariableNamingRule","preserve");
nframes = A0.frame(end);

%% Pre-filtering of localizations (local density, then spot brightness)
%  limits the number of spots (nmax) to avoid long computation times
nmax = 2e6; 
if(height(A0)>nmax)
    % First filter based on local density
    radius = 3*median(A0.("uncertainty [nm]")); 
    n_min = 20;
    [~,d]=knnsearch(A0{:,3:4},A0{:,3:4},'K',n_min + 5);
    d(:,1)=[]; % Delete first column of 0's
    n_neighbors = sum(d < radius,2);
    A = A0(n_neighbors>n_min, :); % Write to new Matrix A
    fprintf('\nUsing %d density filtered localizations out of %d for DME, representing %f %% of data \n', ...
        height(A), height(A0), (height(A)/height(A0)*100))
    % Second filter based on intensity if necessary
    intensity = A.("intensity [photon]");
    if (length(intensity) > nmax)
        prc = 100*nmax/length(intensity);
        thr = prctile(intensity,100-prc);
        A = A(intensity>thr,:);
        fprintf('Limiting dataset to %g spots with highest intensity', nmax);
    end
else
    A = A0;
end

%% Convert coordinates and uncertainties to pixel units
pixelsize = 103.5; % 103.5 nm
locs_px = (1/pixelsize)*A{:,3:4}; % convert nm to pixel units
framenums = A.frame;
frameskips = setdiff(1:nframes, framenums);
if length(frameskips) > 5
    warning('more than 5 frames are without localizations');
end
% Use ThunderSTORM's uncertainty estimates
if any("uncertainty [nm]" == string(A.Properties.VariableNames)) 
    unc_px = A.("uncertainty [nm]")*(1/pixelsize);
    crlb = [unc_px, unc_px];
    fprintf("\nWorking on %s using reported uncertainties \n", name);
else % fixed uncertainty
    fprintf("\nWorking on %s using fixed uncertainty of 0.2px\n", name);
    loc_error = [0.2,0.2]; 
    crlb = repmat(loc_error, length(locs_px), 1);
end
  
%% Set vars for DME algorithm, adapted from Cnossen et al. 2021 (doi.org/10.1364/OE.426620, https://github.com/qnano/drift-estimation, v1.2.1)
% User inputs for RCC drift estimation
usercc = false;    
rcc_timebins = 6;
zoom = 3;
sigma = 1;

% User inputs for DME drift estimation - !keep data types alive!
precision_est = false;                  % Precision estimation (bool)
coarse_frames_per_bin = int32(10);      % Number of bins for coarse est. (int32)
framesperbin = int32(1);                % Number of frames per bin (int32)
maxneighbors_coarse = int32(2000);      % Max neighbors for coarse and precision est. (int32)
maxneighbors_regular = int32(4000);     % Max neighbors for regular est. (int32)
coarseSigma= single(repmat([0.3,0.3], length(locs_px), 1));     % Localization precision for coarse estimation (single/float)                     
max_iter_coarse = int32(200);           % Max iterations coarse est. (int32)
max_iter = int32(1000);                 % Max iterations (int32)
max_iter_est = int32(50);               % Max iterations for precision est (int32)
gradientstep = single(5e-6);            % Gradient (single/float)
usecuda = true;

%% RCC and iterative DME computation
% Compute RCC as initial estimate (usercc=true), or as an independent comparison (usercc = false)
[drift_rcc] = rcc(locs_px, framenums, rcc_timebins, zoom, sigma, 0, usecuda);
if usercc % initial drift estimation by RCC
    rcc0 = drift_rcc; %#ok<UNRCH> 
    fprintf('RCC was used for initial estimation.\n')
else % empty rcc matrix is passed to DME
    rcc0 = repmat([0 0], nframes, 1); 
    fprintf('RCC was NOT used for initial estimation.\n')
end 

% Run iterative DME report run time for parameter tuning
tic 
[drift_dme, d1, d2, d3, drift_set1, drift_set2] = dme_estimate_iterative(locs_px, framenums, crlb, rcc0, usecuda, coarse_frames_per_bin, ...
    framesperbin, maxneighbors_coarse, maxneighbors_regular, coarseSigma, max_iter_coarse,max_iter, gradientstep, precision_est, max_iter_est);
T = toc();
fprintf('\nFirst DME run took %g seconds\n',T)

%% Plot RCC and iterative DME estimates
h = figure;
ndim = 2;
for i = 1:ndim % Plot X and Y drift separately, in nm units
    subplot(ndim,1,i);
    plot(1:1:nframes, drift_dme(:,i)*pixelsize, 'LineWidth', 1);
    hold on
    plot(1:1:nframes, drift_rcc(:,i)*pixelsize, 'LineWidth', 2);
    hold on
    if(exist('d1', 'var')) % Plot DME iterations
        plot(1:1:nframes, d3(:,i)*pixelsize, 'LineWidth', 2);
        hold on
        plot(1:1:nframes, d2(:,i)*pixelsize, 'LineWidth', 2);
        hold on
        plot(1:1:nframes, d1(:,i)*pixelsize, 'LineWidth', 2);
        hold on
    end
    ylim([-200 200])
end

title(name, 'Interpreter', 'none');
legend('Estimated drift (DME)', 'Estimated drift (RCC)', 'Coarse DME 3', 'Coarse DME 2', 'Coarse DME 1') ;
saveas(h, strcat(dir, subfolder, name, '.png'));
saveas(h, strcat(dir, subfolder, name, '.fig'));

%% Apply drift correction to matrix and save
locs_nm0 = A0{:,3:4};
corr_nm0 = locs_nm0 - drift_dme(A0.frame, :)*pixelsize;
A0_corr = A0;
A0_corr(:,3:4) = array2table(corr_nm0);

% Update localization uncertainties 
locprecBefore = NeNA(A0{:,[2 3 4 6]} , name, false);
locprecAfter  = NeNA(A0_corr{:,[2 3 4 6]}, name, false);
ratio = locprecAfter/locprecBefore;
A0_corr.("uncertainty [nm]") = A0.("uncertainty [nm]") * ratio;
fprintf('NeNa Localization precision (frame2frame) improved from %f nm to %f nm\n', locprecBefore, locprecAfter);

% Save corrected localization table
id = strcat(subfolder, 'DME_');
fprintf('Saving results table ... \n');
f_saveLocTable(dir, id, name, ext, A0_corr);

diary off;


%% ------------------- Functions -------------------
% DME function, adapted from Cnossen et al. 2021 (doi.org/10.1364/OE.426620, https://github.com/qnano/drift-estimation, v1.2.1)
function [drift, driftC1, driftC2, driftC3, drift_set1, drift_set2] = dme_estimate_iterative(position, framenumbers, crlb_initial, drift_guess, usecuda, coarse_frames_per_bin, ...
    framesperbin, maxneighbors_coarse,maxneighbors_regular, coarseSigma, max_iter_coarse,max_iter, gradientstep, precision_est, max_iter_est)
    
    % initialize variables
    pos = single(position);
    crlb = single(crlb_initial);
    framenum = int32(framenumbers);
    drift = single(drift_guess);
    numspots = int32(size(pos,1));
    maxdrift = single(0); 
    
    % Compute sets for precision estimate
    % New method to split data in a zebra pattern 
    [~, split_axis] = max(var(pos(:,1:2), 0, 1)); % find axis of greater variance
    binsizepx = 20; % divide axis in 20 px intervals
    set1 = sin(pi/binsizepx*pos(:,split_axis)) > 0; 
    set2 = ~set1;
    
    % Scale factors applied to 'coarse_frames_per_bin' in the first two coarse iterations
    fac1 = 500;
    fac2 = 50;
    
    % Factor applied to 'gradientstep' in the first two coarse iterations
    stepfactor = 10; 
    % Step size in the final estimation
    finalstep = single(1e-6);

    % EXTRA iterative steps
    % Coarse estimation 1, bin size 5000 frames
    fprintf('Computing coarse drift estimation (1)...\n');
    [driftC1, ~, ~] = call_dme_lib('dme', usecuda, pos, coarseSigma*4, framenum, numspots, drift, ...
        coarse_frames_per_bin*fac1,gradientstep*stepfactor,maxdrift,maxneighbors_coarse, max_iter_coarse);
    if(anynan(driftC1))
        error("DME optimization failed in coarse1.");
    end
    
    % Coarse estimation 2, bin size 500 frames
    fprintf('Computing coarse drift estimation (2)...\n');
    [driftC2, ~, ~] = call_dme_lib('dme', usecuda, pos, coarseSigma*2, framenum, numspots, driftC1, ...
        coarse_frames_per_bin*fac2, gradientstep*stepfactor, maxdrift, maxneighbors_coarse, max_iter_coarse);
    if(anynan(driftC2))
        error("DME optimization failed in coarse2.");
    end
    
    % Main routine
    % Coarse estimation final, bin size 10 frames
    fprintf('Computing coarse drift estimation (3)...\n');
    [driftC3, ~, ~] = call_dme_lib('dme', usecuda, pos, coarseSigma, framenum, numspots, driftC2, ...
        coarse_frames_per_bin,gradientstep, maxdrift, maxneighbors_coarse, max_iter_coarse);
    if(anynan(driftC3))
        error("DME optimization failed in coarse3.");
    end
    
    % Regular drift estimation, bin size 1 frame
    fprintf('Computing final drift estimation...\n');
    [drift, ~, ~] = call_dme_lib('dme', usecuda, pos, crlb, framenum, numspots, driftC3, ...
        framesperbin,finalstep, maxdrift, maxneighbors_regular, max_iter);
    if(anynan(drift))
        error("DME optimization failed in final step.");
    end

    % Precision estimate, optional (computation intensive)
    % Computes two independent estimates after splitting the dataset, then computes RMSE
    if precision_est
        fprintf('Computing precision set 1 drift estimation...\n');
        [drift_set1, ~, ~] = call_dme_lib('dme', usecuda, pos(set1, :), crlb(set1,:), framenum(set1), int32(size(pos(set1, :),1)), drift, framesperbin,gradientstep ...
        ,maxdrift,maxneighbors_regular, max_iter_est);
        fprintf('Computing precision set 2 drift estimation...\n');
        [drift_set2, ~, ~] = call_dme_lib('dme', usecuda, pos(set2, :), crlb(set2,:), framenum(set2), int32(size(pos(set2, :),1)), drift, framesperbin,gradientstep ...
        ,maxdrift,maxneighbors_regular, max_iter_est);
        
        % Save drift sets for plotting
        drift_set1 = drift_set1 - mean(drift_set1);
        drift_set2 = drift_set2 - mean(drift_set2);
        L = min(length(drift_set1), length(drift_set2));
        diff = drift_set1(1:L,:)-drift_set2(1:L,:);
        rmsd = sqrt(mean(diff.^2));
        fprintf(2, ['\nRMSD of drift traces on split dataset = ', num2str(rmsd)] )
    else 
        drift_set1 = NaN;
        drift_set2 = NaN;
    end
    
    % Create output drift traces centered on 0/0
    drift = drift - mean(drift);
    driftC1 = driftC1 - mean(driftC1);
    driftC2 = driftC2 - mean(driftC2);
    driftC3 = driftC3 - mean(driftC3);
end

% NeNa algorithm adapted from Martens et al 2021 (https://doi.org/10.1007/s00418-014-1192-3) Module 9b
function locprec = NeNA(data, id, do_plot)
    % take 5% of data in regular intervals
    frames = mod(data(:,1), 2000) >= 1900;
    data = data(frames, :);
    
    % Nearest-neighbour distance calculation
    maxDistance = 120;
    tracked_data = [data zeros(size(data,1),2)];
    tracksCounter = 1;
    for fr = 0:max(tracked_data(:,1))-1
        [tracked_data,tracksCounter] = commonFunctions.creating_trajectories(tracked_data,fr,fr+1,maxDistance,tracksCounter);
    end
    
    % NeNA localization precision determination
    jds = tracked_data(tracked_data(:,6)>0,6);
    if(do_plot)    % currently giving error
        figure
    else
        figure('visible','off');
    end

    % We plot a histogram of next-frame nearest neighbor distances
    h1 = histogram(jds, 150, 'Normalization','Probability','DisplayName','Nearest Neighbour distance');
    x = h1.BinEdges(2:end);
    x = x-x(1)/2;
    ft = fittype('(r./(2.*a.*a)).*exp((-1).*r.*r./(4.*a.*a)).*A + (F/(w*sqrt(pi/2))) .* exp(-2.*((r-rc)./w).*((r-rc)./w)) + O.*r','independent','r');
    NeNAfit = fit(x', h1.Values', ft, 'StartPoint', [0.5, 0.5, 1, 15, 15, 100]);
    locprec = NeNAfit.a;
    
    hold on
    plot(x,NeNAfit(x),'DisplayName',['NeNA fit: ' num2str(NeNAfit.a) ' nm'],'LineWidth',3)
    legend()
    title(strcat('NeNA fit of ', id), 'Interpreter', 'none')
end

% Function to save localization table
function f_saveLocTable(dir, id, name, ext, A0)
    savename = strcat(dir, id, name, ext);
    for x = 1:10
        if isfile(savename)
            savename = strcat(dir, id, name, '_', num2str(x), ext);
        else
            break;
        end
    end
    writetable(A0,savename);
    fprintf('saved as: %s \n', savename);
end
```
