## Supplementary Software for "Nanoscale architecture and dynamics of Ca_V_1.3 channel clusters in cardiac myocytes revealed by single channel nanoscopy": DNA_PAINT_2_MolecularMapping.m

```
% Input: Reconstruction image of merged localizations (.tif) 
%   rendered as 5 nm gaussians on 4 nm pixel grid using ThunderSTORM
%   (these parameters assume localization precision sigma ~ 4 nm)
% Output: Molecular map based on H-maxima (stored in variable: 'points')
%   annotated image (green = accepted, red = rejected) and polygonal ROI saved to file

%% File processing
clearvars
close all
[filename,folder] = uigetfile('*.tif','Select a file');
file = strcat(folder, '\', filename);
fprintf('working on: %s\n', file)

%% Set parameters and call main analysis function
pxsize = 4; % nm
% these parameters need to be set depending on the image intensity scale and background localization density 
% applying a local density filter in ThunderSTORM is recommended
% H-maxima parameter (detection sensitivity)
h = 0.002;
% intensity threshold to exclude spots
thr = 0.15;

% Main function, returns molecular map in µm units
points = Hmaxima(file, pxsize, h, thr); 

% Use the resulting molecular map 'points' for subsequent cluster analysis routine (DBSCAN, NND)
%DNA_PAINT_3_ClusterAnalysis


%% Main analysis function to detect H-maxima (molecular positions)
function [points] = Hmaxima(file, pxsize, h, thr)
    roifile = file + ".mat";
    [A, ~] = imread(file);

    % Show blurred image for ROI selection
    figure
    imshow(imgaussfilt(A,10),DisplayRange=[0 thr]);
    hold on
    colormap("hot")
    
    %% Load or define Region of Interest
    if(exist(roifile))
        f_ui = uifigure;
        uialert(f_ui,'Click OK after defining the ROI in the image','Roi is set.', 'CloseFcn', @(h,e)resume(f_ui), Modal=false, icon='info');
        load(roifile);
        fprintf('Roi was loaded from file, modify and press any key to continue...\n');
        roi.Parent = gca;
        newROI = images.roi.Polygon(gca);
        newROI.Position = ROI.Position;
        ROI = newROI;
        drawnow
        disp('entering pause')
        uiwait(f_ui)
    else
        msgbox('Please create a polygonal ROI in the image (cell boundary)', createmode="modal");
        disp('Please create a polygonal ROI in the image (cell boundary)')
        ROI = drawpolygon;
    end
    save(roifile, 'ROI');
    fprintf('ROI was saved to file.\n');
    
    % Detect local maxima
    A_filt = imhmax(A,h); 
    A_maxima = imregionalmax(A_filt);
    
    %% Report weighted centroid positions and maxima intensity
    spots = regionprops("table",A_maxima,A,"WeightedCentroid", "MaxIntensity");
    idxvalid = (spots.MaxIntensity > thr) & inROI(ROI,spots.WeightedCentroid(:,1),spots.WeightedCentroid(:,2));
    spots_valid = spots(idxvalid,:);
    spots_invalid = spots(~idxvalid,:);
    fprintf('Found %i maxima in the image, %f%% in ROI and over threshold', height(spots), 100*height(spots_valid)/height(spots))
    
    % Render the image and zoom in
    figure
    imshow(A, DisplayRange=[0 3*thr])
    colormap("hot")
    centerpoint = mean(spots_valid.WeightedCentroid,1);
    roisize = 1000; % pixels
    axis([centerpoint(1)-(roisize/2), centerpoint(1)+(roisize/2), centerpoint(2)-(roisize/2), centerpoint(2)+(roisize/2)])
    title("H-maxima over background")
    hold on
    
    % Draw accepted (green) and rejected (red) spots as an overlay
    viscircles(spots_valid.WeightedCentroid,ones(size(spots_valid,1),1)*4,'Color','g', EnhanceVisibility=0, LineWidth=1);
    viscircles(spots_invalid.WeightedCentroid,ones(size(spots_invalid,1),1)*4,'Color','r', EnhanceVisibility=0, LineWidth=1);
    points = spots_valid.WeightedCentroid * pxsize / 1000;

    function resume(fig)
        disp('program resumed');
        uiresume
        close(fig)
    end

end
```
