## Supplementary Software for "Nanoscale architecture and dynamics of Ca_V_1.3 channel clusters in cardiac myocytes revealed by single channel nanoscopy": DNA_PAINT_3_ClusterAnalysis.m

```
% Input: Point set representing a molecular map (variable 'points', unit µm)
% Output: DBSCAN clusters, NNDs, Cluster analysis, cumulative results file

%% Set parameters
eps = 100;
showMap = true;
showFig = true;
saveResults = true;

%% Run DBSCAN clustering algorithm
% with parameters epsilon = 100 nm, minpts = 3
idx = dbscan(points, eps/1000, 3);

% Separate points into clustered and non-clustered array
idxNCL = idx(idx == -1);
idxCL = idx(idx >= 0);
pointsNCL = points(idx == -1, :);
pointsCL = points(idx >= 0, :);

%% Plot clustering results
h = figure;
axis equal
if(~showMap)
    h.Visible = false;
end
set(gca,'XAxisLocation','top','YAxisLocation','left','ydir','reverse');
colors = turbo(numel(idxCL));
colors = colors(randperm(length(colors)), :);
hold on
axis equal
legend('off')
grid('minor');
set(gca,'xtick',0:1:100)
set(gca,'ytick',0:1:100)

% Draw molecule positions as 10 nm circles 
radius = 0.005; 
viscircles(pointsNCL,radius*ones(size(pointsNCL,1),1), Color=[0.5 0.5 0.5], EnhanceVisibility=0);
viscircles(pointsCL,radius*ones(size(pointsCL,1),1), Color='k', EnhanceVisibility=1);

%% Cluster quantification
% Preallocate arrays
num_cl = max(idx);
areas = zeros(num_cl,1);
area_ismin = zeros(num_cl,1);
molcount = zeros(num_cl,1);
clusters = cell(num_cl,1);
pointsCL_NND1=zeros(numel(idxCL), 1);
pointsCL_N3ND=zeros(numel(idxCL), 1);

% Get NNDs between all points
[points_NND1, points_N3ND] = getNND(points, 3);

% Get NNDs between unclustered points
[pointsNCL_NND1, pointsNCL_N3ND] = getNND(pointsNCL, 3);

% get total area via boundary function (can be concave)
[bound, A_bound] = boundary(pointsCL);

e = 0; % keep track of errors
errortext = '';
warning('off','MATLAB:polyshape:repairedBySimplify')

for i = 1:num_cl
    cluster = points(idx == i,:);
    clusters{i} = cluster;
    try
        [bound, ~] = boundary(cluster);
        col = colors(i,:);
        buffer = 10; % 10 nm buffer radius = 5nm physical + 5nm uncertainty
        polyout = polybuffer(polyshape(cluster(bound,:)), buffer/1000);
        plot(polyout, 'FaceColor', col, 'FaceAlpha', 0.2);
        A = polyout.area;
    catch exception
        errortext = getReport(exception,'basic');
        e = e + 1;
        A = NaN;
    end
    molcount(i) = size(cluster,1);
    areas(i) = A;
end
if(e>0)
    fprintf('%i X error: %s',e,errortext)
end

% GetNND function returns NND1 and NxND for each point in a set
[nnd1, n3nd] = getNND(pointsCL, 3);
pointsCL_NND1 = nnd1;
pointsCL_N3ND = n3nd;


%% Compute descriptive statistics
diams = 1000*2*sqrt(areas/pi);
dens = molcount./areas;
dens(dens == Inf) = 0;
cl_fract = 100*numel(idxCL)/numel(idx);
cl_dens = num_cl/A_bound;

mean_count = mean(molcount);
mean_area = mean(areas);
mean_diams = mean(diams);
mean_dens = mean(dens);

med_count = median(molcount);
med_area = median(areas);
med_diams = median(diams);
med_dens = median(dens);

med_nnd1 = median(pointsCL_NND1);
med_n3nd = median(pointsCL_N3ND);

% Report statistics to command window
fprintf('\n-----\nFound %i clusters containing %i points\n', num_cl, numel(idxCL));
fprintf('Clustered fraction = %f %% \nCluster density = %f/µm² \nMean MolCount = %f\nMean Cl. Diameter = %f\nMean Mol.Dens = %f\n', ...
    cl_fract, cl_dens, mean_count, mean_diams, mean_dens);
fprintf('Median NND1 = %f | Median NND3avg = %f\n', med_nnd1, med_n3nd);

%% Save result tables (will append to existing tables)
if(saveResults)
    id = convertCharsToStrings(file);
    stats = table(id, eps, num_cl, cl_fract, cl_dens, mean_count, mean_area, mean_diams, mean_dens, med_nnd1, med_n3nd, med_count, med_area, med_diams, med_dens); %#ok<UNRCH> 
    results = table(repelem(id,num_cl)', molcount, areas, diams, dens);
    savedir = folder + "PooledResults\";
    if(~exist(savedir, 'dir'))
        mkdir(savedir);
    end
    % save summary statistics
    writetable(stats,savedir + 'stats_summary.xls','WriteMode','append')
    % save single cluster data
    writetable(results,savedir + 'stats_clusters.xls','WriteMode','append');
    % save nnd values
    writematrix(points_NND1,savedir + 'points_NND1.csv','WriteMode','append')
    writematrix(pointsCL_NND1,savedir + 'pointsCL_NND1.csv','WriteMode','append')
end

%% Plot histograms
if(showFig)
    figure %#ok<UNRCH> 
    subplot(2,4,1)
        histogram(points_NND1, BinWidth=10)
        xlim([0 300])
        title('NND1 for all channels')
    subplot(2,4,2)
        histogram(pointsCL_NND1, BinWidth=10);
        title('NND1 for clustered channels')
        xlim([0 300])
    subplot(2,4,3)
        histogram(points_N3ND, BinWidth=10);
        xlim([0 300])
        title('N3ND for all channels')    
    subplot(2,4,4)
        histogram(pointsCL_N3ND, BinWidth=10);
        xlim([0 300])
        title('N3ND for clustered channels')
    subplot(2,4,5)
        histogram(molcount, 0.5:1:30);
        title('Cluster channel count')
    subplot(2,4,6)
        histogram(areas, BinWidth=0.004);
        title('Cluster area')
    subplot(2,4,7)
        histogram(dens, BinWidth=200);
        title('Intra-cluster channel density')
    subplot(2,4,8)
        scatter(areas, molcount, 25, 'MarkerFaceColor','b','MarkerEdgeColor','b','MarkerFaceAlpha',.05,'MarkerEdgeAlpha',.05)
        xlim([0 0.05]) 
        ylim([0 40])
        title('Channel count over cluster area')
end

%% NND analysis function for 1st and x-th nearest neighbor
function [nnd1, nXnd] = getNND(points, X) 
    [~,d_knn]=knnsearch(points, points,'K',X+1); % e.g. input 4 = 3 closest nnds + 0-value
    d_knn(:,1)=[]; % delete first column of 0's
    nnd1 = 1000*d_knn(:,1);
    nXnd = 1000*mean(d_knn,2);
end
```
