## Supplementary Software for "Nanoscale architecture and dynamics of Ca_V_1.3 channel clusters in cardiac myocytes revealed by single channel nanoscopy": ImageJ_1 STED CaV cluster analysis.ijm

```
/* ImageJ Macro for automated segmentation of low SNR, spot-like fluorescent signals within a defined size range
 * this method can deal well with high signal intensity variation, low SNR and variable local background (as opposed to threshold-based methods)
 * the parameters were optimized for STED images of Halo-Cav1.3 clusters in the basal plasma membrane
 * Input image: unprocessed STED image with annotated cell outline, can be multi-channel
 * Requires the H-Watershed plugin for ImageJ: https://github.com/mpicbg-scicomp/Interactive-H-Watershed | https://imagej.net/plugins/interactive-watershed 
 */

// General parameters for processing
chSTED = 2;
closeAll = true;
saveResults = true;
saveFilteredImg = false;
overlayLineProfiles = true; // draw and save line profile overlay with r2 indication

// Type of bandpass filtering: 1 = FFT band pass; 2 = DoG (Difference of Gaussians); 3 = 1px gaussian + 20px rolling ball 
bandpassType = 1; 

// Background parameter for Part 1: H-watershed segmentation
bgThr = 0; // 0 = histogram-based auto thresholding

// Segmentation parameters for Part 2: H-watershed segmentation and filtering
minSize = 5; // area (px) used to filter watershed ROIs
hMin = 2; // higher H-value adds more region merging and maxima noise resistance
fillPct = 50; // 50 = fwhm
batch = true; // speeds up processing

// Set resolution-based bandpass parameters and calibration-derived single dye brightness
// lowpass:  2.5 px * 25 nm = 62.5 nm minimum object (cluster) size 
// highpass:  20 px * 25 nm = 500 nm  maximum object (cluster) size 
getPixelSize(unit, pixelWidth, pixelHeight);
px_size = round(pixelWidth*1000);

if(px_size == 25 || px_size == 30){
	lowpass = 2.5; 
	// set single dye brightness (SDB) from DNA Origami calibration
	SDB = 32.5;
}else {
	exit("error: unexpected pixel size");
}
highpass = 20; 

// File processing
if(bitDepth() != 16) // applies to raw Abberior files, tif-format conversion is recommended
	exit("Macro aborted: The image is not raw data");
title = getTitle();
id_original = getImageID();
Stack.setDisplayMode("color");
filepath = getDirectory("image");
File.makeDirectory(filepath + "processed\\");
cellroi = filepath + "processed\\CellROI_"+ title + ".roi";
print("Spot analysis for image: " + title + " in filepath: " + filepath);

if(selectionType == -1){
	print("There is no preselected cell ROI. Loading a saved Cell ROI.");
	open(cellroi);
	if(File.exists(cellroi)==false)
		 exit("There is no selection");		
}

if(filepath=="" || saveResults == false){
	print("Using the temp folder for testing");
	filepath = getDirectory("temp");
}

// Pre-processing of the STED channel 
Stack.setChannel(chSTED); 
Overlay.remove();
run("Fire");
setMinAndMax(0, 10 + SDB);

run("Set Measurements...", "display redirect=None decimal=6");
run("Measure"); // create results row with file name
CellArea = getValue("Area"); // get cell area to calculate particle density
setResult("CellArea", nResults-1, CellArea);

// Convert STED channel image to 32-bit for bandpass
run("Select None");
roiManager("reset");
run("Duplicate...", "ignore"); 
setOption("ScaleConversions", false); // no value rescaling
run("32-bit");

// Bandpass filtering
// Option 1: apply FFT band pass filter, correct for offset 
if(bandpassType == 1){
	run("Bandpass Filter...", "filter_large=" + highpass  + " filter_small=" + lowpass  + " suppress=None tolerance=5");
	getStatistics(area, mean, min, max, sd, histogram);
	run("Subtract...", "value=" + mean +" slice");
	rename(title + " - FFT band pass");
	setMinAndMax(0, SDB);
}
// Option 2: apply DoG filter
if(bandpassType == 2){
	min_sigma = lowpass / 2.355;
	max_sigma = highpass / 2.355;
	rename("DoG_bg");
	id_bg = getImageID();
	run("Duplicate...", "title=DoG_signal");
	id_signal = getImageID();
	run("Gaussian Blur...", "sigma=" + min_sigma);
	selectImage(id_bg);
	run("Gaussian Blur...", "sigma=" + max_sigma);
	//run("Square Root", "slice"); // take square root of background to save peak intensities and prevent halo effects
	imageCalculator("Subtract 32-bit", id_signal, id_bg);
	// rename filter output and close partial outputs
	rename(title + " - DoG filtered");
	close("DoG_*");
}
// Option 3: gaussian filter + rolling ball bg subtraction (less effective for low SNR/ amplitude)
if(bandpassType == 3){
	run("Gaussian Blur...", "sigma=1 stack");
	run("Subtract Background...", "rolling=20 sliding disable stack");
	run("Min...", "value=0"); // ignore negative values
	
}

id_filtered = getImageID();

// Clear signals outside of cell ROI
run("Restore Selection");
print("Saving: " + cellroi);
saveAs("Selection", cellroi);
run("Clear Outside", "stack");


// --------------------------------------------------------------------------------------
// --- PART 1: AUTOMATIC THRESHOLDING ---
// ----------------------------------------------------------------------------------------
// Simple method: mean+std+offset (not used)
getStatistics(area, mean, min, max, std, histogram);
autoThrSimple = mean + std;
print("Mean+SD = " + autoThrSimple);

// Advanced method: 90th histogram percentile
if(bgThr == 0){
	getHistogram(values, counts, 1000); //only works on 32 bit (filtered) image
	cumsum = 0;
	total = getValue("Area raw");
	limit = 0.9*total;
	for (i=0; i<values.length; i++){
	  cumsum += counts[i];
	  if (cumsum > limit){//getHeight()*getWidth()){
	  	histo = values[i];
	  	i = values.length; //ends loop
	  }
	}
	print("histogram 90th percentile: " + histo);

	bgThr = 2 + 0.5*histo;
	print("Auto threshold: " + bgThr);
}
setMinAndMax(0, 5*bgThr); 
run("Select None");


// --------------------------------------------------------------------------------------
// --- PART 2: H-WATERSHED SEGMENTATION AND FILTERING ---
// ----------------------------------------------------------------------------------------
// Run H-Watershed and iterate over label map to add to ROI manager
run("H_Watershed", "impin=["+getTitle()+"] hmin="+hMin+" thresh="+bgThr+" peakflooding="+fillPct+" outputmask=false allowsplitting=true");
if(batch){
	setBatchMode("hide");
}
max = getValue("Max");
for(i=max;i>0;i--){
	setThreshold(i,i);
	run("Create Selection");
	if(selectionType() == 9){
		//split discontinous ROIs produced by watershed
		roiManager("Split"); 
	}else{ 
		roiManager("Add");
	}
}

if(closeAll) 
	close(); //close watershed image

// Spot quality filtering by size and intensity
selectImage(id_filtered);
OutS = 0; // size filtering counter
OutI = 0; // int. filtering counter

for (n=0; n<roiManager("count"); ++n) { 
	roiManager("Select", n);
	RoiSize = getValue("Area raw");
	Mean = getValue("Mean");
	if(RoiSize < minSize){
		// Out by minsize
		roiManager("delete");
		n-=1;
		OutS+=1;
	}else{
		if (Mean < 1.5*bgThr){ // requires mean to be at least 50% over background
			// Out by mean intensity
			roiManager("delete");
			n-=1;
			OutI+=1;
		}else{
			// Spot has passed filtering
			roiManager("rename", "Spot " + n);
		}
	}
}
nspots = roiManager("count");
print("Final spot count: " + nspots + " after filtering out: "+ OutI +" by intensity and "+ OutS + " by minsize");

if(batch)
	setBatchMode("exit and display");

// Save all spot ROIs
roiManager("deselect");
roipath = filepath + "processed\\ROIset_" + title + ".zip";
roiManager("save", roipath);

// Save filtered image
roiManager("show all without labels");
if(saveFilteredImg){
	saveAs("tif", filepath + "processed\\Filt_" + title);
}
roiManager("deselect");
if(closeAll) 
	close();
	
	
// --------------------------------------------------------------------------------------
// --- PART 3: CLUSTER ANALYSIS ---
// ----------------------------------------------------------------------------------------
// Iterate over spots and measure, later create summary statistics
// Initialize arrays for spot measurement
ArArea = newArray(nspots);
ArCircsize = newArray(nspots);
ArRaw_RawIntDen = newArray(nspots);
ArRaw_Mean = newArray(nspots);
ArRaw_MeanBackground = newArray(nspots);
ArRaw_IntDenFinal = newArray(nspots);
ArRoundness = newArray(nspots);
ArMaximum = newArray(nspots);
ArX = newArray(nspots);
ArY = newArray(nspots);
ArFWHM = newArray(nspots);
ArMolC = newArray(nspots);
ArMolD = newArray(nspots);
print("Measuring spots");

// switch to raw image for measurements
selectImage(id_original);
if(batch)
	setBatchMode("hide");
for (n=0; n<nspots; ++n) {
	roiManager("Select", n);
	ArArea[n] = getValue("Area");
	ArCircsize[n] = 2000*sqrt(getValue("Area")/PI);
	offset = 0.5; // pixel shift in coordinate system
	ArX[n] = (getValue("XM") / px_size * 1000) - offset;
	ArY[n] = (getValue("YM") / px_size * 1000) - offset;
}
for (n=0; n<nspots; ++n) {
	roiManager("Select", n);
	ArRaw_RawIntDen[n] = getValue("RawIntDen");
	ArRaw_Mean[n] = getValue("Mean");
	ArRoundness[n] = getValue("Round");
	ArMaximum[n] = getValue("Max");
	ArFWHM[n] = fwhm(ArX[n],ArY[n],0.85); // drawing takes a long time if displayed
}
if(batch)
	setBatchMode("exit and display");
	
// save raw image with ROI overlay
if(saveResults){
	roiManager("show all without labels");
	saveAs("tif", filepath + "processed\\" + title); 
}

// Local background correction of intensity values
print("Performing bg correction");
if(closeAll)
	setBatchMode(true); // does not create a visible image of bg subtraction
run("Select None");
run("Duplicate...", "ignore");
run("32-bit"); // set 32-bit to enable NaN values

// Delete cluster signals with a 2 pixel margin, then create a donut-shaped ROI for background measurement
for (n=0; n<nspots; ++n) {
	roiManager("Select", n);
	run("Enlarge...", "enlarge=2 pixel");
	run("Set...", "value=NaN");
	run("Enlarge...", "enlarge=2 pixel");
	roiManager("update")
}

// Perform array computations
for (n=0; n<nspots; ++n) {
	roiManager("Select", n);
	ArRaw_MeanBackground[n] = getValue("Mean");
	// Correction for background intensity resulting in final raw intensity
	ArRaw_IntDenFinal[n] = ArRaw_RawIntDen[n]*(1-(ArRaw_MeanBackground[n]/ArRaw_Mean[n])) ;
	// report maxima and signal density
	ArMaximum[n] = ArMaximum[n] - ArRaw_MeanBackground[n];
	ArMolC[n] =  ArRaw_IntDenFinal[n] / SDB;
	ArMolD[n] =  ArMolC[n] / ArArea[n];
}
if(closeAll) 
	close();
setBatchMode(false);

// Create table outputs and save
printArrays(); // function call
File.makeDirectory(filepath + "processed\\");
Table.save(filepath + "processed\\ClusterResults_" + title + ".csv");		

// Discard background ROIs and reload cluster ROIs
selectImage(id_original);
roiManager("reset"); 
roiManager("open", roipath);

// Save log file
selectWindow("Log");
saveAs("text", filepath + "processed\\Log " + title);

print("Task completed.");
beep();


// --------------------------------------------------------------------------------------
// --- Print Array function ---
// ----------------------------------------------------------------------------------------
function printArrays(){
	// report summary statistics
	setResult("nspots", nResults-1, nspots);
	setResult("Threshold", nResults-1, bgThr);
	setResult("SpotDens", nResults-1, nspots/CellArea);
	
	Array.getStatistics(ArArea, min, max, mean, stdDev);
	setResult("Cl. Area", nResults-1, mean);
	Array.getStatistics(ArCircsize, min, max, mean, stdDev);
	setResult("Cl. Diam", nResults-1, mean);
	Array.getStatistics(ArMolC, min, max, mean, stdDev);
	setResult("MolCount", nResults-1, mean);
	Array.getStatistics(ArMolD, min, max, mean, stdDev);
	setResult("MolDens", nResults-1, mean);
	Array.getStatistics(ArRaw_IntDenFinal, min, max, mean, stdDev);
	setResult("IntDen_final", nResults-1, mean);
	Array.getStatistics(ArRaw_RawIntDen, min, max, mean, stdDev);
	setResult("IntDen_raw", nResults-1, mean);
	Array.getStatistics(ArRaw_MeanBackground, min, max, mean, stdDev);
	setResult("MeanBg", nResults-1, mean);
	Array.getStatistics(ArRoundness, min, max, mean, stdDev);
	setResult("Roundness", nResults-1, mean);
	
	// get mean FWHM, ignorin invalid data
	meanFWHM = 0;
	count = 0;
	nval = 0;
	for (n=0; n<nspots; ++n) {
		val = ArFWHM[n];
		if(val > 0){ // not NaN
			meanFWHM += val;
			nval += 1;
		}
	}
	setResult("FWHM", nResults-1, meanFWHM/nval);
	
	// report single cluster data in another table
	Table.create("Single spot measurements");	
	Table.setColumn("spot num", Array.getSequence(nspots));
	Table.setColumn("posX", ArX);
	Table.setColumn("posY", ArY);
	Table.setColumn("Area", ArArea);
	Table.setColumn("Circize", ArCircsize);
	Table.setColumn("MolCount", ArMolC);
	Table.setColumn("MolDens", ArMolD);
	Table.setColumn("B_final", ArRaw_IntDenFinal);
	Table.setColumn("B_rawID", ArRaw_RawIntDen);
	Table.setColumn("MeanSig", ArRaw_Mean);
	Table.setColumn("MeanBg", ArRaw_MeanBackground);
	Table.setColumn("Roundness", ArRoundness);
	Table.setColumn("MaxOverBg", ArMaximum);
	Table.setColumn("FWHM", ArFWHM);
}


// --------------------------------------------------------------------------------------
// --- FWHM line profiles function ---
// ----------------------------------------------------------------------------------------
function fwhm(px,py,gof){
	linewidth = 3; // iteratively increased
	lineSize = 10;
	
	spotMin = 2*px_size; //nyquist
	spotMax = highpass*px_size; 
	lineSize=lineSize*0.5; 
	if(overlayLineProfiles){
		run("Overlay Options...", "set");
		makePoint(px, py, "add");
	}
	
	// Horizontal line profile
	lw = 0;
	do{	
		lw++;
		makeLine(px-lineSize,py,px+lineSize,py,lw); 

		Yval=getProfile(); 
		Xval=newArray(Yval.length); 
		for(j=0;j<Yval.length;j++){
			Xval[j]=j*px_size/1000; 
			if(Yval[j] == 0) Yval[j] = NaN;
		} 
		Fit.doFit("Gaussian", Xval, Yval); 
		h_r2=Fit.rSquared;
		h_d_sigma=Fit.p(3); 
		h_FWHM=1000*abs(2*sqrt(2*log(2))*h_d_sigma);
		if(h_FWHM <= spotMin || h_FWHM >= spotMax)
			h_r2 = 0;
	}while(h_r2 < gof && lw < linewidth);
	if(overlayLineProfiles){
		run("Overlay Options...", "stroke=green width=0.1 fill=none set");
		run("Add Selection...");	        
	}

	horOK = h_r2 > gof;
	
	// Vertical line profile
	lw = 0;
	do{	
		lw++;
		makeLine(px,py-lineSize,px,py+lineSize, lw); 
		Yval=getProfile(); 
		Xval=newArray(Yval.length);		
		for(j=0;j<Yval.length;j++){ 
			Xval[j]=j*px_size/1000; 
			if(Yval[j] == 0) Yval[j] = NaN;
		}
		Fit.doFit("Gaussian", Xval, Yval); 
		v_r2=Fit.rSquared; 
		v_d_sigma=Fit.p(3); 
		v_FWHM=1000*abs(2*sqrt(2*log(2))*v_d_sigma);
		if(v_FWHM <= spotMin || v_FWHM >= spotMax)
			v_r2 = 0;
	}while(v_r2 < gof && lw < linewidth);
	if(overlayLineProfiles){
		run("Overlay Options...", "stroke=green width=0.1 fill=none set");
		run("Add Selection...");	        
	}

	vertOK = v_r2 > gof;
	run("Select None"); 
	
	// Evaluate fits
	if(horOK && vertOK){
		if(overlayLineProfiles){
			makeRectangle(px-lineSize+0.5,py-lineSize+0.5, lineSize*2, lineSize*2); 
			run("Overlay Options...", "stroke=green width=0.25 fill=none set");
			run("Add Selection...");
		}	
		if(h_r2 > v_r2){
			return h_FWHM;
		}else{
			return v_FWHM;
		}
		

	}else{ 
		if(horOK == false && vertOK == false){
			// no good fit
			if(overlayLineProfiles){
				makeRectangle(px-lineSize+0.5,py-lineSize+0.5, lineSize*2, lineSize*2); 
				run("Overlay Options...", "stroke=red width=0.25 fill=none set");
				run("Add Selection...");
			}
			return NaN;
		}else { 
			// only one good fit
			if(overlayLineProfiles){
				makeRectangle(px-lineSize+0.5,py-lineSize+0.5, lineSize*2, lineSize*2); 
				run("Overlay Options...", "stroke=yellow width=0.25 fill=none set");
				run("Add Selection...");	
			}
			if(horOK){
				return h_FWHM;
			}
			if(vertOK){
				return v_FWHM;
			}
	
		}
	}  
}
```
