## Supplementary Software for "Nanoscale architecture and dynamics of Ca_V_1.3 channel clusters in cardiac myocytes revealed by single channel nanoscopy": ImageJ_2 Confocal CRU colocalization.ijm

```
// ImageJ Macro to compute three-channel segmentation and quantify colocalization of CaV signal-mass to other channels 
// Input image: 3-channel unprocessed confocal image with annotated cell outline

// Parameter setup by user, define channel order
saveResults = true;
ChCaV = 1;
ChRYR = 2;
ChJPH = 3;

// File processing
filepath = getDirectory("image");
title = getTitle();
cellroi = filepath + "processed\\CellROI_"+ title + ".roi";
File.makeDirectory(filepath + "processed\\");
roipath = filepath + "processed\\ROI_" + title + "channel_";
idOrig = getImageID();

// load cell outline ROI or save it (if drawn)
if(selectionType == -1){
	open(cellroi);
}else{
	saveAs("Selection", cellroi);
}

// convert image to floating-point format (lossless, no rescale)
run("Crop");
run("Duplicate...", "duplicate");
roiManager("reset");
run("Select None");
run("Conversions...", " ");
run("32-bit");

// run bandpass filter and correct offset for each channel
run("Bandpass Filter...", "filter_large=40 filter_small=4 suppress=None tolerance=5 process");
Stack.getDimensions(width, height, channels, slices, frames);
for (i = 1; i <= channels; i++) {
	Stack.setChannel(i);
	getStatistics(area, mean, min, max, sd, histogram);
	run("Subtract...", "value=" + mean +" slice");
}

// remove negative values and convert to 8-bit for thresholding
Stack.setChannel(1);
run("Min...", "value=0 stack");
run("Multiply...", "value=3 stack"); // prevent quantization at low signal levels
run("8-bit");
run("Restore Selection");
Stack.setDisplayMode("composite");

// enhance contrast (only for display)
Stack.getDimensions(width, height, channels, slices, frames);
for (i = 1; i <= channels; i++) {
	Stack.setChannel(i);
	run("Enhance Contrast...", "saturated=1");
}

// save bandpass version
saveAs("tif", filepath + "processed\\bp_" + title); 
idFilt = getImageID();
run("Duplicate...", "duplicate");

// apply auto thresholding to all channels
run("Auto Local Threshold", "method=Otsu radius=20 parameter_1=0 parameter_2=0 white stack");
idSeg = getImageID();


// run Analyze Particles for each channel and save ROIs
setBatchMode(true);
for (i = 1; i <= channels; i++) {
	selectImage(idSeg);
	Stack.setChannel(i);
	run("Duplicate...", " "); // isolate single channel image
	setThreshold(1, 255);
	run("Analyze Particles...", "size=8-Infinity pixel show=Nothing add");
	roiManager("deselect");
	roiManager("combine"); // create union roi of cluster signals
	roiManager("reset"); // discard the invididual cluster rois
	Roi.setName("Ch" + i);
	roiManager("add");
	roiManager("save", roipath + i + ".zip");
	roiManager("reset");
	close();
}
setBatchMode(false);

// Save binarized image
run("Select None");
roiManager("show none");
saveAs("tif", filepath + "processed\\thr_" + title); 

// Load segmentation of channels in original (image channel) order
roiManager("open", roipath + "1.zip"); 
roiManager("open", roipath + "2.zip");
roiManager("open", roipath + "3.zip");

// Processing of binary overlaps (intersection) in defined order
roiManager("Select", newArray(ChCaV-1,ChRYR-1));
roiManager("and");
Roi.setName("CaV AND RYR");
roiManager("add");

roiManager("Select", newArray(ChCaV-1,ChJPH-1));
roiManager("and");
Roi.setName("CaV AND JPH");
roiManager("add");

roiManager("Select", newArray(0,1,2));
roiManager("and");
Roi.setName("CaV AND RYR AND JPH");
roiManager("add");

selectImage(idOrig);
Stack.setChannel(ChCaV);

// Measure the signal mass of CaV1.3 for different binary maps
count = roiManager("count");
run("Set Measurements...", "area display redirect=None decimal=6");
roiManager("select", ChCaV-1);
run("Measure"); // create measurement row
bCaV = measureIntDen("CaV"); // measure all Cav1.3 signal above threhold
roiManager("select", count-3);
bCaV_RYR = measureIntDen("CaV-RYR"); // measure above-threshold Cav1.3 signal + threshold RYR
roiManager("select", count-2);
bCaV_JPH = measureIntDen("CaV-JPH"); // measure above-threshold Cav1.3 signal + threshold JPH2
roiManager("select", count-1);
bCaV_triple = measureIntDen("triple"); // measure above-threshold Cav1.3 signal + threshold RyR2 + threshold JPH2

// Compute colocalization based on Manders M1
setResult("M1-RYR", nResults-1, bCaV_RYR/bCaV);
setResult("M1-JPH", nResults-1, bCaV_JPH/bCaV);
setResult("M1-triple", nResults-1, bCaV_triple/bCaV);

// Measure the area of different binary maps, not considering signal intensity
roiManager("select", ChCaV-1);
measureArea("Ch1 all"); 
roiManager("select", count-3);
measureArea("CaV-RYR"); 
roiManager("select", count-2);
measureArea("CaV-JPH");
roiManager("select", count-1);
measureArea("triple");

// Functions for table output
function measureArea(label){
	setResult("Area " + label, nResults-1, getValue("Area"));
}

function measureIntDen(label){
	val = getValue("IntDen");
	setResult("IntDen " + label, nResults-1, val);
	return val;
}
```
