## Supplementary Software for "Nanoscale architecture and dynamics of Ca_V_1.3 channel clusters in cardiac myocytes revealed by single channel nanoscopy": ImageJ_3 Confocal CTT cluster thresholding.ijm

```
// ImageJ Macro to perform thresholding-based cluster segmentation and quantify the resulting particles 
// Input image: Unprocessed confocal image with annotated cell outline, can be multi-channel

// Set the channel to be analyzed
Stack.setChannel(4);

// File processing
filepath = getDirectory("image");
title = getTitle();
roi_cell = filepath + "processed\\CellROI_"+ title + ".roi";
File.makeDirectory(filepath + "processed\\");
idOrig = getImageID();

// load cell outline ROI or save it (if drawn)
if(selectionType == -1){
	open(roi_cell);
}else{
	saveAs("Selection", roi_cell);
}

// Isolate cluster channel and clear outside
run("Select None");
run("Duplicate...", " "); // single channel
run("Gaussian Blur...", "sigma=1 stack");
run("Restore Selection");
run("Clear Outside");

// convert to 8-bit (no rescaling) for thresholding and measure total brightness
run("Conversions...", "calibrate");
run("8-bit");
run("Enhance Contrast...", "saturated=0.5");
getMinAndMax(min, max);

// Get whole-cell intensity measurement
run("Set Measurements...", "area mean standard min integrated display redirect=None decimal=6");
run("Measure"); 

// Cluster segmentation by auto-thresholding
run("Auto Threshold", "method=Moments ignore_black show white setthreshold"); // Otsu needs "ignore_black"
run("Analyze Particles...", "size=6-120 Infinity pixel exclude summarize overlay composite");
run("Select None");
setMinAndMax(min, max);
run("Overlay Options...", "stroke=none width=0 fill=none set");

// Save processed image with cluster annotation
saveAs("tif", filepath + "processed\\thr_" + title); 
selectImage(idOrig);
close();

// Results and Summary Table must be saved after all images have been processed
```
