## Supplementary Software for "Nanoscale architecture and dynamics of Ca_V_1.3 channel clusters in cardiac myocytes revealed by single channel nanoscopy": SPT_1_Diffusion_analysis_trackit.m

```
% Input: Single particle tracks and MSD values ('batch' and 'analysis' file, grouped by cells/recordings) 
% --> generated by TrackIt software (Kuhn et al. 2021, doi.org/10.1038/s41598-021-88802-7)
% Output: Quantitative diffusion and confinement analysis in various figures and console outputs

%% File and variable processing
close all
clearvars
pxsize = 0.1035;  % µm
frametime = 0.03; % s

% Load batch data file and analysis data file generated by TrackIt ('Export to Matlab workspace', include all movies)
load('JF646_trackit_batch.mat');
load('JF646 analysis.mat')
Results = vertcat(batch(:).results);
Tracks.All = [Results(:).tracks]';
Tracks.JD = vertcat(Results(:).jumpDistances);
nTracks = numel(Tracks.All);

% Preallocate arrays
Tracks.Int = cell(nTracks,1);
Tracks.IntMean = zeros(nTracks,1);
Tracks.length = vertcat(Results(:).trackLengths);

% Iterate over all tracks
for i = 1:nTracks
    track = Tracks.All{i}; 
    intensity = track(1:end-1,4); % discard last point
    Tracks.Int{i} = intensity;
    Tracks.IntMean(i) = mean(intensity);
    l = min(5, floor(size(intensity,1)/2));
    Tracks.lengthLocs(i,1) = size(track,1);
    Tracks.lengthSecs(i,1) = (track(end,1)-track(1,1))*frametime;
end

%% Plot track length + intensity histogram
figure
subplot(2,2,1)
histogram(Tracks.length, [0:5:100 Inf], 'Normalization','probability');
xlabel('track length (frames)')
ylabel('rel. frequency')
xlim([0 105])

subplot(2,2,2)
histogram(Tracks.lengthLocs, [0:2:100 Inf], 'Normalization','probability');
xlabel('track length (localizations)')
ylabel('rel. frequency')
xlim([0 105])

subplot(2,2,3)
histogram(Tracks.IntMean, [0:100:2000 Inf], 'Normalization','probability');
xlabel('Mean spot intensity of track')
ylabel('rel. frequency')
xlim([0 2100])

%% Retrieve MSD and Mean jump distance values
D_MSD.cells = allAnalysisResults.msdDiffConst;
D_MSD.all = vertcat(D_MSD.cells{:});
MJD.cells = allAnalysisResults.meanJumpDists;
MJD.all = vertcat(MJD.cells{:});
D_MSD.cellMed = cellfun(@median, D_MSD.cells);
fprintf('Mean D = %f, Median D = %f \n', mean(D_MSD.all), median(D_MSD.all));
fprintf('Median D across cells: %f ± %f (Mean ± SD) \n', mean(D_MSD.cellMed), std(D_MSD.cellMed))
assert(numel(Tracks.IntMean) == numel(D_MSD.all));

%% Compute confinement statistics
% Preallocate arrays
rgyr = nan(nTracks,1);
rgyr_300ms = nan(nTracks,1);
rgyr_1s = nan(nTracks,1);
chull_A_300ms = nan(nTracks,1);
chull_A_first = nan(nTracks,1);
chull_d_300ms = nan(nTracks,1);

% Compute confinement values for tracks with 10 or more localizations
i_filt = find(Tracks.lengthLocs > 10);
for i = i_filt' 
    track = Tracks.All{i}; % discard last point
    frames = track(:,1);
    track_um = pxsize * track(:,2:3);
    
    % Interpolate gap frames
    track_interp = interp1(frames,track_um,frames(1):frames(end),'spline');
    
    % Compute convex hull within sliding window
    a = ConvHullSliding(track_interp, 10);
    chull_A_300ms(i,1) = a; % convex hull area, over sliding window
    chull_d_300ms(i,1) = 1000 * 2 * sqrt(a/pi); % equivalent diameter
    
    % Compute gyration radius within sliding window
    rgyr(i) = 1000 * RadiusGyr(track_interp);
    rgyr_300ms(i,1) = 1000 * RadiusGyrSliding(track_interp,10);
    rgyr_1s(i,1) = 1000 * RadiusGyrSliding(track_interp,33);
end

% Report statistics to console window
fprintf('%f %% of tracks have more than 10 locs and are used for confinement analysis:\n  ', ...
    100*numel(i_filt)/nTracks);

fprintf(['Convex hull area (300 ms) = %f ± %f \n  ' ...
    'Convex hull diameter (300 ms) = %f ± %f \n  ', ...
    'Radius of gyration (300 ms) = %f ± %f \n  '], ...
    mean(chull_A_300ms, 'omitnan'), std(chull_A_300ms, 'omitnan'), ...
    mean(chull_d_300ms, 'omitnan'), std(chull_d_300ms, 'omitnan'), ...
    mean(rgyr_300ms(:,1), 'omitnan'), std(rgyr_300ms(:,1), 'omitnan'));

%% Plot confinement analysis histograms
figure
subplot(2,2,1) % shown in Figure 3
histogram(chull_A_300ms, 0:0.0016:0.04, 'Normalization','probability');
xlabel({'A(conv. hull), 300 ms (µm²)'});%;'tracks ≥ 10 locs'})
ylabel('rel. frequency')

subplot(2,2,2)
histogram(chull_d_300ms, 0:20:300, 'Normalization','probability');
xlabel({'d(conv. hull), 300 ms (nm)';'tracks ≥ 10 locs'})
ylabel('rel. frequency')

subplot(2,2,3) % shown in Figure 3
histogram(rgyr_300ms, 0:8:200, 'Normalization','probability');
xlabel({'r(gyration), 300 ms (nm)'});%;'tracks ≥ 10 locs'})
ylabel('rel. frequency')

subplot(2,2,4)
histogram(rgyr_1s, 0:10:200, 'Normalization','probability');
xlabel({'r(gyration), 1 s (nm)';'tracks ≥ 10 locs'})
ylabel('rel. frequency')

%% Plot jump distances and diffusion coefficients
figure
subplot(2,2,1)
histogram(1000*MJD.all, 1000*[0:0.01:0.3 Inf], 'Normalization','probability');
xlabel('Mean jump distance (nm)')
ylabel('rel. frequency')
xlim([0 310])

subplot(2,2,2)
histogram(D_MSD.all, [0:0.001:0.03 Inf], 'Normalization','probability');
xlabel('D_{fit} (µm²/s)')
ylabel('rel. frequency')
xlim([0 0.031])

% log10 histograms of diffusion coefficients
subplot(2,2,3)
x = logspace(-5,0,21);
histogram(D_MSD.all, [10^(-10) x 10] ,'Normalization','probability');
set(gca,'xscale','log')
xlabel('D_{fit} (µm²/s)')
ylabel('rel. frequency')
xlim([10^(-5.25) 10^(0.25)])
xticks(logspace(-5,0,6))

subplot(2,2,4)
histogram(log10(D_MSD.all), [-10 -5:0.25:0],'Normalization','probability');
xticks(-5:1:0)
xlim([-5.25 0])
xlabel('log_{10}(D_{fit}) (µm²/s)')
ylabel('probability density')

%% Generate scatter plots to check variable correlations as potential bias
figure
subplot(1,3,1)
scatter(Tracks.IntMean, D_MSD.all, 8, 'filled', MarkerFaceAlpha=0.2)
xlabel('Mean spot intensity')
ylabel('D_{fit} (µm²/s)')
set(gca,'yscale','log')
xlim([0 2000])

subplot(1,3,2)
scatter(Tracks.length, D_MSD.all, 8, 'filled', MarkerFaceAlpha=0.2)
xlabel('Track length')
ylabel('D_{fit} (µm²/s)')
set(gca,'yscale','log')
xlim([0 200])

subplot(1,3,3)
scatter(Tracks.length, Tracks.IntMean, 8, 'filled', MarkerFaceAlpha=0.2)
xlabel('Track length')
ylabel('Mean spot intensity')
xlim([0 200])
ylim([0 2000])
set(gcf,'Position',[100 100 900 225])

% Compute and report immobile fractions
D_MSD.cells_mean = cellfun(@mean,D_MSD.cells);
D_MSD.cells_n = cellfun(@length,D_MSD.cells);
D_MSD.cells_n_start = cumsum([1; D_MSD.cells_n]);

Fract_immobile=@(in1) 100*sum(in1 < 1e-3)/numel(in1);
immobile = Fract_immobile(D_MSD.all);
immobile_cell = cellfun(Fract_immobile, D_MSD.cells);

fprintf('Immobile fraction pooled = %.1f %% (Mean, defined as D < 0.001)\n', mean(immobile))
fprintf('Immobile fraction per cell = %.1f ± %.1f %% (Mean ± SD, defined as D < 0.001)\n\n', mean(immobile_cell), std(immobile_cell))


%% ------------------- Functions -------------------
% Simple radius of gyration across whole trajectory
function [rms_dist] = RadiusGyr(locs) 
    center = mean(locs,1);
    delta  = center - locs; 
    rms_dist = rms(vecnorm(delta'));
end

% Radius of gyration within a sliding window
function [rms_dist] = RadiusGyrSliding(locs, window) 
    if(size(locs,1) < window)
        rms_dist = nan;
        return
    end
    % get center point within sliding window
    center =movmean(locs,window,1); 
    % get distance to center point
    delta  = center - locs; 
     % discard start and end overhangs
    buffer = floor(window/2);
    delta = delta(1+buffer:end-buffer,:);
    % get rms of distance to center point (moving window gyration radius)
    rms_dist = rms(vecnorm(delta')); 
end

% Convex hull area within a sliding window
function [a_mean] = ConvHullSliding(locs, window) 
    n = size(locs,1);
    if(n < window)
        a_mean = nan;
        return
    end
    i = 0;
    sum = 0;
    while(i + window <= n)
        [~,a] = convhull(locs(i+1:i+window,:));
        sum = sum + a;
        i = i + 1;
    end
    a_mean = sum / i;
end
```
