## Supplementary Software for "Nanoscale architecture and dynamics of Ca_V_1.3 channel clusters in cardiac myocytes revealed by single channel nanoscopy": SPT_2_SimulationImmobileLocError.m

```
% Input: Experimental track length distribution, sigma estimate
% Output: Estimates of apparent MJD and RGyr caused by the localization error
% Estimated run-time: 10 min for 10,000 Mone-Carlo runs X 5,000 tracks X 25 localizations

clearvars
%% Choose between loading experimental track lengths or setting fixed values
% Load experimental track length distribution
% load('JF646_trackit_batch.mat');
% Results = vertcat(batch(:).results);
% nTracks = size([Results(:).tracks],2);
% lengths = vertcat(Results(:).trackLengths);

% Alternative: Use a constant track length and number:
nTracks = 5000;
lengths = ones(nTracks,1)*25;

%% Setup parameters
% Set sigma = estimated localization precision in nm
sigma_mean = 25;
sigma_sdev = 5;  
sigma_min = 1; 
sigma_min_count = 0;

% Initiate Monte-Carlo variables
nruns = 1000;
montecarlo_mjd_mean = 0;
montecarlo_mjd_sd = 0;
montecarlo_rgyr_mean = 0;
montecarlo_rgyr_sd = 0;

%% Run simulation
for montecarlo = 1:nruns
    % Generate a sigma value for each track
    sigma = normrnd(sigma_mean, sigma_sdev, nTracks, 1);
    sigma = max(sigma, sigma_min);
    mjd = nan(nTracks,1);
    rgyr = nan(nTracks,1);
    
    % Simulate and quantify all tracks
    for i = 1:nTracks
        % Generate random localizations around (0/0)
        xy = normrnd(0, sigma(i), lengths(i), 2);
        % Get frame-to-frame displacements
        jd = diff(xy);
        % Calculate Mean Jump Distance
        dist = vecnorm(jd');
        mjd(i) = mean(dist);
        % Calculate Radius of Gyration
        rgyr(i) = RadiusGyr(xy);
        if(sigma(i) == sigma_min)
            sigma_min_count = sigma_min_count + 1;
        end
    end
    montecarlo_mjd_mean  = montecarlo_mjd_mean  + mean(mjd);
    montecarlo_mjd_sd    = montecarlo_mjd_sd    + std(mjd);
    montecarlo_rgyr_mean = montecarlo_rgyr_mean + mean(rgyr);
    montecarlo_rgyr_sd   = montecarlo_rgyr_sd   + std(rgyr);
end

% Determine and report Monte-Carlo results
montecarlo_mjd_mean  = montecarlo_mjd_mean  / nruns;
montecarlo_mjd_sd    = montecarlo_mjd_sd    / nruns;
montecarlo_rgyr_mean = montecarlo_rgyr_mean / nruns;
montecarlo_rgyr_sd   = montecarlo_rgyr_sd   / nruns;

fprintf(['Monte-Carlo simulation with nruns = %i resulted in \n' ...
    'MJD = %f ± %f nm and RGyr = %f ± %f nm (average Mean ± average SD)\n'], ...
    nruns, montecarlo_mjd_mean, montecarlo_mjd_sd, montecarlo_rgyr_mean, montecarlo_rgyr_sd);


%% Function to calculate radius of gyration
% A sliding window is not necessary, since the position is by definition immobile
function [rms_dist] = RadiusGyr(locs) 
    center = mean(locs,1);
    delta  = center - locs; 
    rms_dist = rms(vecnorm(delta'));
end
```
